# Crowdsourced Protein Design: Lessons From the Adaptyv EGFR Binder Competition

**DOI:** 10.1101/2025.04.17.648362

**Authors:** Tudor-Stefan Cotet, Igor Krawczuk, Filippo Stocco, Noelia Ferruz, Anthony Gitter, Yoichi Kurumida, Lucas de Almeida Machado, Francesco Paesani, Cianna N. Calia, Chance A. Challacombe, Nikhil Haas, Ahmad Qamar, Bruno E. Correia, Martin Pacesa, Lennart Nickel, Kartic Subr, Leonardo V. Castorina, Maxwell J. Campbell, Constance Ferragu, Patrick Kidger, Logan Hallee, Christopher W. Wood, Michael J. Stam, Tadas Kluonis, Süleyman Mert Ünal, Elian Belot, Alexander Naka, Adaptyv Competition Organizers

**Affiliations:** Adaptyv; Centre for Genomic Regulation, Pompeu Fabra University; Department of Biostatistics and Medical Informatics, University of Wisconsin–Madison; Morgridge Institute for Research; School of Frontier Engineering, Kitasato University; Instituto Oswaldo Cruz, Fiocruz; Department of Chemistry and Biochemistry, University of California San Diego; BioLM; École Polytechnique Fédérale de Lausanne; School of Informatics, University of Edinburgh; Hearth Industries; Cradle Bio; Synthyra; Center for Bioinformatics & Computational Biology, University of Delaware; School of Biological Sciences, University of Edinburgh; Unaffiliated; Science Corporation

## Abstract

In this report, we summarize and analyze the 2024 Adaptyv protein design competition. Participants used computational and Machine Learning (ML) methods of their choice to design proteins that bind the Epidermal Growth Factor Receptor (EGFR), a key drug target involved in cell growth, differentiation, and cancer development.

Over 1,800 designs were submitted across two rounds. Of these, 601 proteins were selected and characterized for expression and binding affinity to EGFR, with competitors both optimizing existing binders (*K*_*D*_ = 1.21 nM) and creating *de novo* binders (*K*_*D*_ = 82 nM).

All selected designs were experimentally validated using Adaptyv’s automated Bio-Layer Interferometry (BLI) pipeline. This competition illustrates the potential of crowdsourcing to drive creativity and innovation in protein design. However, it also exposed key challenges, such as the lack of standardized benchmarks, experimental design targets, and robust computational metrics for method comparison. We anticipate that future competitions will address these gaps and further motivate progress in computational protein design.

## 1 Introduction

Computational protein design holds immense potential to address complex challenges across healthcare, agriculture, sustainability, and biotechnology. Initially, protein design approaches leveraged physics-based energy functions and simulations to engineer novel protein structures, exemplified by the design of the first *de novo* protein fold Top7 in 2003 [Kuh+03]. Another significant milestone was achieved with the design of the first *de novo* protein binder in 2011 [Fle+11], demonstrating the feasibility of computationally engineering proteins to interact specifically with targeted molecular surfaces. Despite these early successes, traditional protein design methodologies faced considerable limitations, including computational inefficiencies and difficulties in reliably predicting protein structure and properties. Significant breakthroughs in the field were enabled by the integration of ML in the 2000s, Deep Learning (DL) in the 2010s, and pre-trained foundation models in the early 2020s. A prototypical example of this transformation is AlphaFold2 (AF2), which enabled highly accurate structural predictions of both natural and *de novo* designed proteins [Jum+21; Eva+22], substantially accelerating protein design campaigns and improving success rates both *in silico* and experimentally [Ben+23]. Importantly, the open-source release of AF2 significantly accelerated the development of novel computational methods and profoundly impacted the trajectory and pace of innovation within the protein design field.

The impact of DL networks on the success rate of designs of valid protein structures and sequences has enabled a shift in focus towards designing functional proteins, including enzymes, biosensors, and specifically tailored protein binders [CLH24]. Specifically, efforts are being invested into improving the design of binders. Numerous studies demonstrate the potential applications of *de novo* designed binders in therapeutics, diagnostics, and fundamental biological research [Che+17; Cao+20; Joh+24; Tor+25]. Traditionally, binder design involves selecting a suitable target binding site, docking compatible protein backbone scaffolds, and subsequently designing sequences that stabilize both the interface and the overall folded structure [Cao+22; Ben+23]. This multi-objective optimization makes binder design particularly complex, requiring simultaneous optimization for proper protein folding, interface stability, and specificity. This is further complicated by the influence of environmental binding conditions, the inherent subtleties of structural changes that influence binding interactions, and the ongoing limitations of predictive computational models. However, the use of DL methods has enabled models to implicitly capture and optimize some of these properties, significantly increasing the success rates of designing successful binders and greatly reducing the scale of experimental screening required [Wat+23; Ben+23].

Despite rapid advancements, assessing and comparing new design methods remains challenging due to the absence of standardized benchmarks, consistent *in silico* evaluation metrics, and publicly available, validated target datasets. Many open questions remain about how diverse and often competing design goals (expression yield, stability, immunogenicity, potency, etc.) can be effectively reconciled during design or addressed during model training. Community-driven initiatives and open competitions are playing a crucial role towards addressing these challenges. Some noteworthy examples which have fostered collaboration, stimulated innovation, and generated rich datasets that are publicly accessible are: Adaptyv’s EGFR Binder Competition, Liberum Bio’s Protein Design Winter Games, the BioML Challenge by UT Austin, Biomod 2024, the Open Protein Design Tournament [Arm+24], the NeurIPS 2024 Leash BELKA Kaggle challenge, and the ongoing AIntibody challenge [Era+24]. Beyond benchmarking state-of-the-art computational methods, these competitions also democratize access to experimental validation, enabling researchers worldwide to test and refine their design approaches. Consequently, such open challenges catalyse methodological advances and promote transparency, reproducibility, and community engagement within the field.

In this context, the 2024 Adaptyv EGFR binder competition provided a unique platform to evaluate a diverse array of computationally designed proteins, ranging from *de novo* designed miniproteins to antibody formats, in a standardized experimental setting. Participants utilized various ML-guided methods and protein engineering strategies to tackle the formidable challenge of designing binders against EGFR, a clinically relevant and highly challenging therapeutic target [Tyd+24]. Here, we explore the outcomes of the competition, analyze experimental insights, and discuss key learnings with implications for the broader protein design community.

## 2 Competition Overview

In July 2024, Adaptyv organized this competition to design binders against EGFR, a clinically relevant target often associated with unchecked cell growth in cancerous tumours [UMY21]. The goal was to create an open competition for biology with equivalent standards of repeatability to those seen the broader ML and Data Science fields, such as with Kaggle competitions [Don23]. To this end, Adaptyv aimed to contribute open benchmarks and drive the development of well-understood metrics, methods, and standard workflows to assist in rationalizing computational protein engineering.

The challenge was to create a new binder for EGFR, distinct from known therapeutics and with the greatest affinity possible. EGFR was chosen for this challenge based on its clinical significance and its well-defined neutralization pathways amenable to *de novo* binder design. Successful therapies like Cetuximab indicate that directly blocking EGFR‘s ligand binding site is sufficient for clinical neutralization [LKF08]. As previous computational efforts successfully produced *de novo* EGFR binders, the competition was designed to be tractable but challenging[Cao+22].

Once announced in June 2024 (Figure 1), participants were invited to submit protein sequences up to a certain length, requiring a minimum number of mutations from known proteins to ensure diversity and relevance of results. Several hundred submitted designs were chosen by Adaptyv for synthesis and affinity characterization using Adaptyv’s automated cell-free protein expression and BLI platform for affinity measurement. Competition winners (top 3 in Round 1 and top 5 in Round

**Figure 1:**
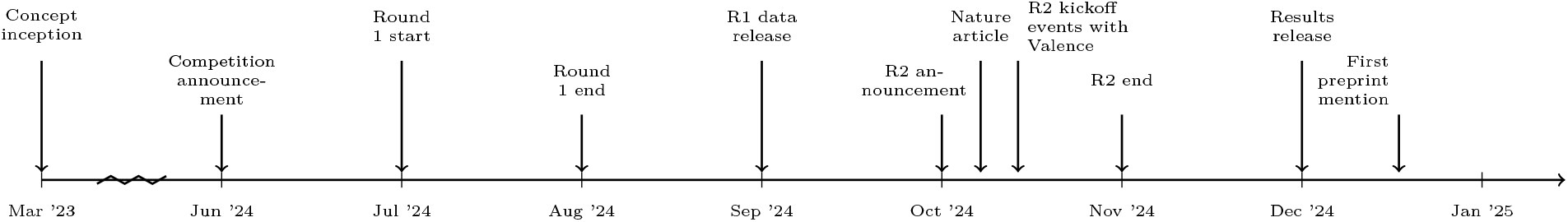
Adaptyv EGFR Competition timeline.

2) were selected each round based on the best *K*_*D*_ achieved, each of them receiving 10 free assays from Adaptyv. Round 1 results were published in September 2024.

Given an overwhelming response, Adaptyv organized a second round of the competition, launched in October 2024. This round was co-organized with Valence Labs and Dimension Capital, which together with Adaptyv hosted physical kick-off events in the US and Europe. It featured sponsored computational resources by the cloud provider Modal, increasing the usability of computational methods; and partial sponsoring of DNA costs by Twist Bioscience, enabling Adaptyv to increase the total number of tested sequences. The length limit of eligible EGFR binders was also extended. Round 2 results were revealed in December 2024.

### 2.1 Submission Eligibility

Crucially, participants were not restricted to commercial or non-commercial/academic institutions, and there was no registration process. Instead, participants simply submitted their designs to a form hosted by Adaptyv, which were automatically scored using the *in silico* scores and added to the daily leaderboard. Each participant was allowed to submit as many sequences as they wanted, but only their top 10 per the *in silico* scores were ranked and displayed in the leaderboard.

The only eligibility requirements were that the sequences must:

1. Be fewer than 200 residues for Round 1 of the Adaptyv Bio EGFR Binder Design Competition (R1); and fewer than 250 for Round 2 of the Adaptyv Bio EGFR Binder Design Competition (R2).
2. Differ in at least 10 residues from publicly known binders (e.g., patented sequences from USPTO).
3. Contain only natural amino acids and be single-chain (e.g., scFv or nanobody for antibodies).

Due to the ranking criteria being based on AlphaFold2 Multimer (AF2-M) metrics (Interface Predicted Aligned Error (iPAE) and for R2 also Interface Predicted Template Modeling Score (ipTM)), an implicit requirement within the competition was that sequences must be foldable by AF2-M [Eva+22], more specifically the ColabFold [Mir+22] implementation using MMSeqs2 for generating the Multiple Sequence Alignment (MSA) [SS17].

### 2.2 Sequence Selection

In R1, sequences were selected by ranking the Interface Predicted Aligned Error (iPAE) from AF2-M structural predictions [Jum+21] [Eva+22]. iPAE roughly represents an error metric between the predicted alignment of EGFR and the binder. This method has previously been used for computationally screening binders [Ben+23].

Following feedback on the ranking metric from R1 and observation of the correlation between other metrics and overall expression rates [Haa24], for R2 a multi-objective metric was chosen. In R2, sequences were selected using a Borda count aggregate of ESM2 pseudo-log-likelihood [Lin+23] (*not* normalized by sequence length), iPAE, and ipTM scores from AlphaFold. ipTM adds a confidence metric with respect to AlphaFold2’s overall prediction, and the incorporation of ESM2 log-probability adds a measure of sequence naturalness, intended to boost the expression rate compared to R1. Arguably, either metric could be biased against (or for) certain *de novo* or underrepresented folds. Moreover, the unnormalized likelihood was biased towards smaller designs.

In both R1 and R2, additional designs that fell outside of the top-ranked sequences were selected by hand. By doing so, Adaptyv aimed to increase diversity and spread across various design methods, architectures, and pipeline strategies, and test novel approaches.

This increase in variance should strengthen the benchmarking of ML models and statistics based on the competition results. In R1, 100 sequences were ranked and chosen along with an additional 101 sequences selected by hand out of 726. In R2, 100 sequences were ranked and an additional 300 sequences were selected manually out of 1131 submissions. In R2, each designer had at least one submission experimentally validated.

### 2.3 Affinity Characterization

Adaptyv’s automated expression, affinity measurement, and post-processing workflow was used to validate the selected designs (Figure 2). This starts with a DNA optimization step to ensure adequate expression of all sequences, followed by a cell-free expression protocol, and BLI for measuring binding affinity. The target protein, in this case EGFR, is introduced at different concentrations to measure both the weak and strong interactions to the immobilized binder.

**Figure 2:**
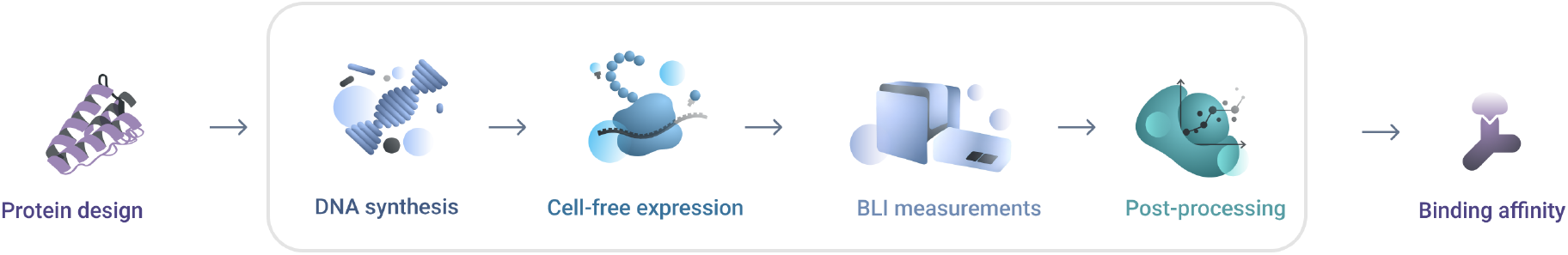
Visual summary of Adaptyv’s workflow for binding affinity measurements.

**Figure 3:**
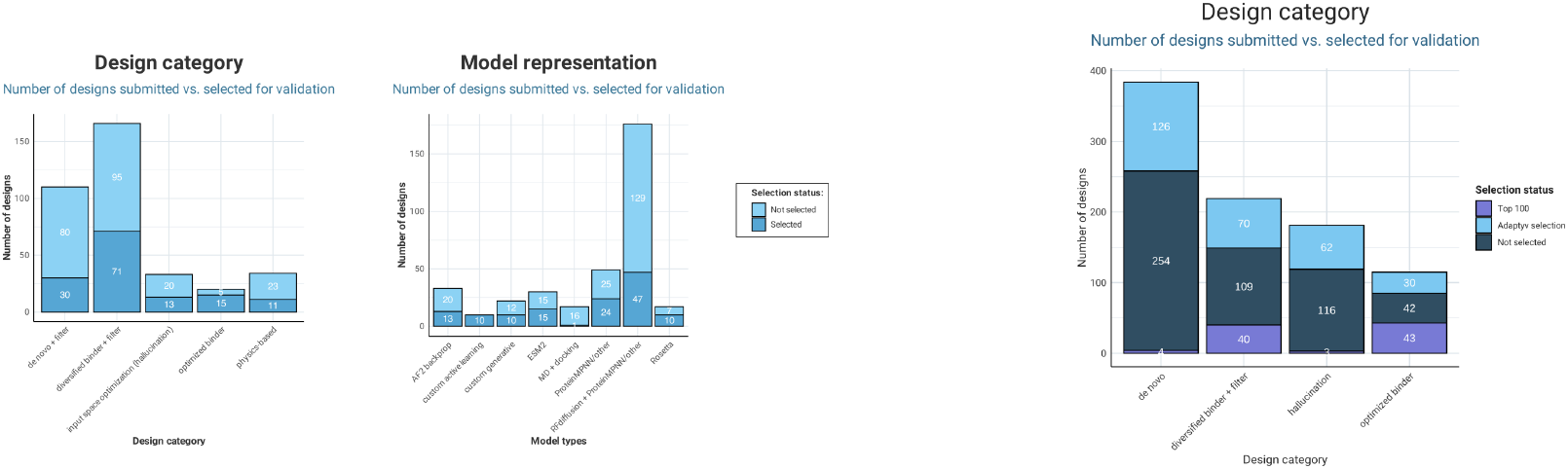
Comparison of the design methods across the two rounds of competition.

**Figure 4:**
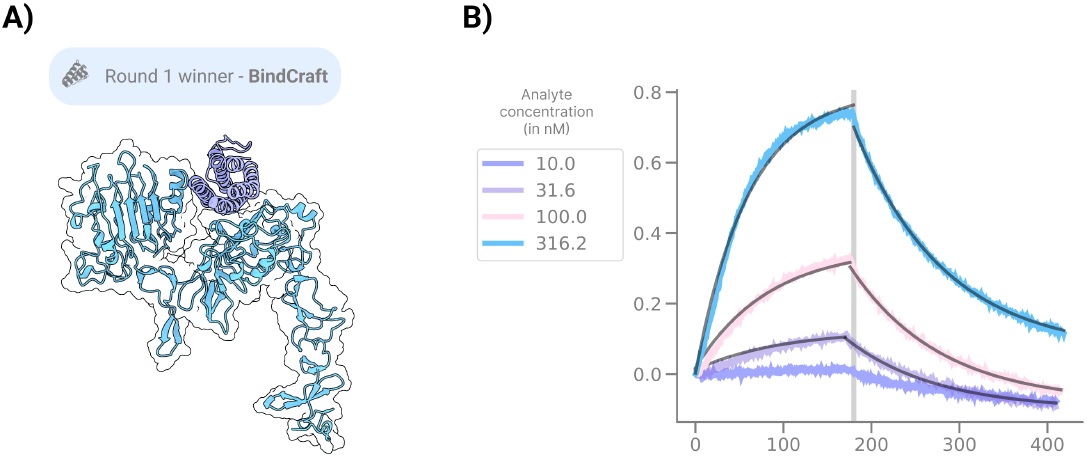
Overview of the R1 winner. (A) Protein structure model of the design bound against EGFR, (B) BLI curve with association/dissociation for different target concentrations for a single replicate.

**Figure 5:**
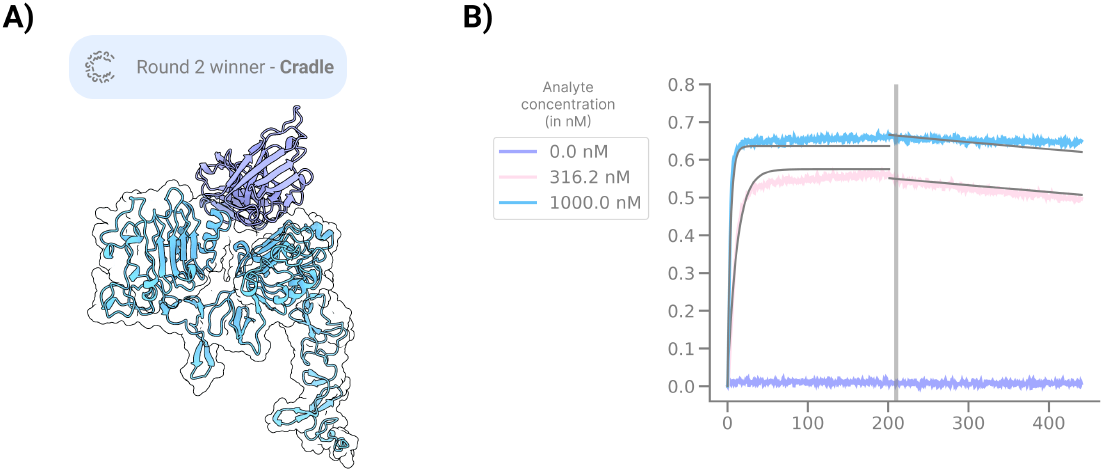
Overview of the R2 winner. (A) Protein structure model of the design bound against EGFR, (B) BLI curve with association/dissociation for different target concentrations for a single replicate.

This is followed by data standardization, a curve-fitting procedure, and quality control steps to ensure that all measurements are reliable [Tea24a]. A minimum of two replicates was run for each design. Additionally, EGF and Cetuximab were run as positive controls. This pipeline was previously validated on IL-7Ra binders obtained from the original RFdiffusion paper [Wat+23], showcasing excellent agreement [Tea24b].

### 2.4 Participation Overview

In the first round, 726 sequences were submitted by approximately 50 designers, including both individual participants and teams. Designers came from around the world, representing diverse backgrounds across industry and academia. Out of these submissions, 146 sequences successfully expressed with 5 novel binders (plus two additional binders later disqualified due to similarity to known sequences). Notably, among participants who provided background information, only about 50% had prior experience in protein design. The 73% expression rate matched typical rates reported in the field, while the binder hit rate was either comparable to or an order of magnitude higher than previous studies [CK24a]. For instance, earlier work on *de novo* EGFR binders reported a 0.01% hit rate, compared to the 2.5% (5 out of 201) observed in R1 [Cao+22].

Subsequent changes to the scoring metrics and to the design strategies [CK24a; CK24b] in R2 were driven by community feedback and made possible through collaborative sharing of expertise within the community. Whether it was these changes, the knowledge sharing spurred by the first round (see section 2.5), or other factors, R2 saw a 95% expression rate (378 of 400 tested) as well as a much higher binding rate with 53 confirmed binders.

Accounting for the larger team sizes (including multiple commercial design teams), R2 more than doubled in the number of participants (130), many of which were first-time designers directly using the tools and learnings that had been shared in the response to R1 [CK24a; CK24b]. Overall, the competition amassed 159 unique participants with 1857 submitted designs.

It is noteworthy for future organizers that despite expecting a last minute rush, in both rounds Adaptyv was surprised by the spike during the last 48 hours, which caused the leaderboard to have delayed updates in R1 and a sizeable shift in sequence distribution to occur in R2. Assuming this pattern will hold, it should be taken as a warning for future organizers to be able to handle at least 10 to 20 times the submissions on the last day than on any other day during the competition.

### 2.5 Post-Competition Response and Highlights

Adaptyv published a series of blog posts accompanying the competition after R1, dubbing them *Protein Optimization* - [CK24c; CK24a; CK24b; CK24d]. Both during and after each round of the competition, the online protein design community was also actively engaged in discussions and sharing ideas. These include various longform posts [Nau25; Cam25; Viz25] and X (Twitter) threads [Ale24] with participants compiling curated collections of social media outputs. Some participants even ordered additional sequences to be tested sharing their results on GitHub and X [Nik24; Git25; Suz25].

The R2 winners (Cradle Bio) have gone into detail on their design methods and results, including organizing webinars and other events to detail their method. Submitting 11 additional designs for testing for their R2 competition prize, they found that all 11 would have won the competition, 3 would have outperformed the original winning design with sub-nanomolar affinity, and the single best design obtained 339 pM. This was whilst maintaining diversity of up to 19 mutations [Fer25].

Other highlights included a Nature news article [Cal24] broadcasting the first round’s results beyond the protein design space, and the fact that within a week of release, the binder data was referenced in academic preprints [Sto+24]. Last but not least, a specific tweet called for the creation of the present manuscript.

## 3 Popular competition protein design methods

### 3.1 Round 1 winning entry: BindCraft

BindCraft [Pac+24] is a computational design pipeline that utilizes AF2-M [Jum+21; Eva+22] backpropagation to generate *de novo* protein binders. The design process is initialized with a random starting sequence, which is then co-folded with the target structure using AF2-M [Eva+22] to produce a predicted complex. This initially suboptimal complex structure guides the improvement of the binder sequence using gradient descent to satisfy multiple design criteria: AF2-M confidence scores, intra- and inter-chain contacts, radius of gyration, and helical content [Gov+24; Dau+22]. Finally, the optimised binder sequence is predicted again in complex with the target using the monomer trained AF2 model [Jum+21] and scored using PyRosetta physics-based metrics [Alf+17].

In the first round of the design competition, authors of the original study used the default settings to generate binders against the beta sheeted interface of EGFR. The top scoring binders according to the AF2 ipTM metric were mostly helical, and five were selected for evaluation. Four of those showed high expression rates and one exhibited the strongest binding affinity in this round of the competition at 491 nM.

In the second round, the authors submitted only beta-sheeted designs from the same campaign, generated by setting the helical loss parameter in BindCraft to −2.0 (default is −0.3). Ten of these designs were tested. Despite the lower overall expression rates observed in R1, all ten showed high expression. One of the designs achieved 82 nM binding affinity to the EGFR target, making it the tightest binding *de novo* binder identified in R2. Overall, 7 out of 8 *de novo* binders in the second round were designed using BindCraft, even though most *de novo* submissions in the competition originated from the RFDiffusion pipeline. Finally, the best R2 BindCraft design had no close structural matches in either the Protein Data Bank (PDB) or AlphaFold Protein Structure Database (AFDB), suggesting that BindCraft can generalize beyond known structural space to produce high-affinity binders.

Other notable BindCraft entries include one participant’s multi-domain submissions each featuring two BindCraft designs with the solubility-enhancement tag GB1 [Hut+97] connected by custom disordered linkers. Five of these composite designs were tested in R2; all five showed high expression but did not bind. Another participant used additional Protein Language Model (pLM) filters [Lin+23; Gel+25] on the BindCraft designs.

### 3.2 Round 2 winning entry: Cradle

The overall winner of the competition (by binding affinity), and the winner of R2, was the ML-guided lead optimization strategy of Cradle [Fer+24], with a binding affinity of 1.21 nM.

As an optimization strategy, a starting point was selected. For this, Cetuximab was used, and then reformatted as a Single-chain Variable Fragment (scFv), joining the variable regions of the light and heavy chains with a glycine-serine linker. This was to meet the competition’s requirements of being single-chain. Mutations were then generated in the framework region. Cetuximab was also used as a positive control in the competition (9.94 nM), allowing for the calculation that this represents an 8.2*×* improvement.

The contestants preserved Cetuximab’s Complementarity-Determining Regions (CDRs), mainly responsible for binding, and focused on mutating the antibody framework regions for improved affinity. The hypothesis was that mutations to the framework region could be targeted to induce small conformational changes in the CDRs, preserving the existing binding mechanism while obtaining carefully controlled improvements [Zho+20]. In contrast, changes directly to the CDRs have the potential to induce far greater structural changes, which could easily prevent or significantly impede binding to EGFR.

The mutations themselves had the potential to be guided by either in-domain data from R1 of the competition, via evolutionary data from public databases, or via *in silico* structure predictions. In this case, the contestants elected to primarily focus on evolutionary data—in particular, since the first round of the competition generated only two weak *de novo* binders. Evolutionarily related sequences were obtained using MMseqs2 [SS17], searching against the uniref30 and colabfold_envdb databases [Mir+22]. Further details are deferred to a future paper, where the contestants aim to describe their work in more detail.

### 3.3 TIMED and DE-STRESS

TIMED is a family of Convolutional Neural Networks (CNNs) for protein sequence design [Cas+24], which uses voxel grids to predict the probability distributions of the 20 amino acids at each position in a sequence. The 10 TIMED designs submitted in R2 of the competition had less than 50% sequence identity with Epidermal Growth Factor (EGF), high expression, and one exhibited *µ*M *K*_*D*_.

To begin with, the contestants analyzed R1 designs using DE-STRESS [SW21] and NetSolP [Thu+21] to identify key factors for expression. DE-STRESS evaluates designed proteins using structural features such as all-atom scoring functions, aggregation propensity, and hydrogen bonding quality. NetSolP predicts solubility and purification usability from ESM sequence embeddings [Riv+21]. This analysis revealed that lower Aggrescan3D aggregation propensity scores [SW21; Kur+19], higher NetSolP usability [Thu+21], and smaller designs were correlated with higher expression. The insights gained from this analysis informed the design process for the sequences submitted in R2.

The contestants redesigned a known EGFR binder, selecting EGF due to its small size, and using the crystal structure of EGF-EGFR (PDB: 1IVO) as the backbone. Sequences were designed using different versions of TIMED: vanilla TIMED, TIMED-Charge (incorporating charge information), Co-TIMED (including EGFR side-chain context), along with DenseCPD and DenseNet [QZ20]. Monte Carlo sampling (T = 0.1) was used to generate 20,000 sequences per model (100,000 in total) and then NetSolP was used to predict the usability of these sequences. Design selection followed two approaches: (1) choosing the top 200 sequences based on NetSolP usability and creating consensus sequences from the top 10 and top quartile, and (2) sampling 20 sequences from the top 1% of NetSolP usability scores. After this, the sequences were folded using ColabFold [Mir+22], and DE-STRESS was used to calculate structural features. Furthermore, the competition metrics (ipTM, iPAE, ESM log-likelihood) were calculated using the Adaptyv GitHub repository^1^, and BUDE FF interaction energy scores with EGFR provided an additional validation step [McI+14].

Finally, the top eight designs from approach (1) were selected based on the metrics described above, along with the two consensus sequences from approach (2), and drMD Molecular Dynamics (MD) simulations were performed to confirm these designs remained stable and properly folded [SNW24].

### 3.4 RFdiffusion

In total, 144 submissions in R1 and 245 submissions in R2 made use of the current State-Of-The-Art (SOTA) RFdiffusion, making it one of the most popular design methods (Figure 6). RFdiffusion [Wat+23] is a diffusion model that was created by fine-tuning the structure prediction model RoseTTAFold [Bae+21] on protein structure denoising tasks. It can perform unconditional generation of proteins, motif scaffolding, and binder design, and can be conditioned on specific binding interface sites and folds. The participants’ use of RFdiffusion in the competition encompassed both fully *de novo* binder generation as well as diversification of existing binders (partial diffusion). This model’s popularity is unsurprising considering its reported experimental hit rate of 19% for *de novo* binders [Wat+23] and its significant accessibility.

**Figure 6:**
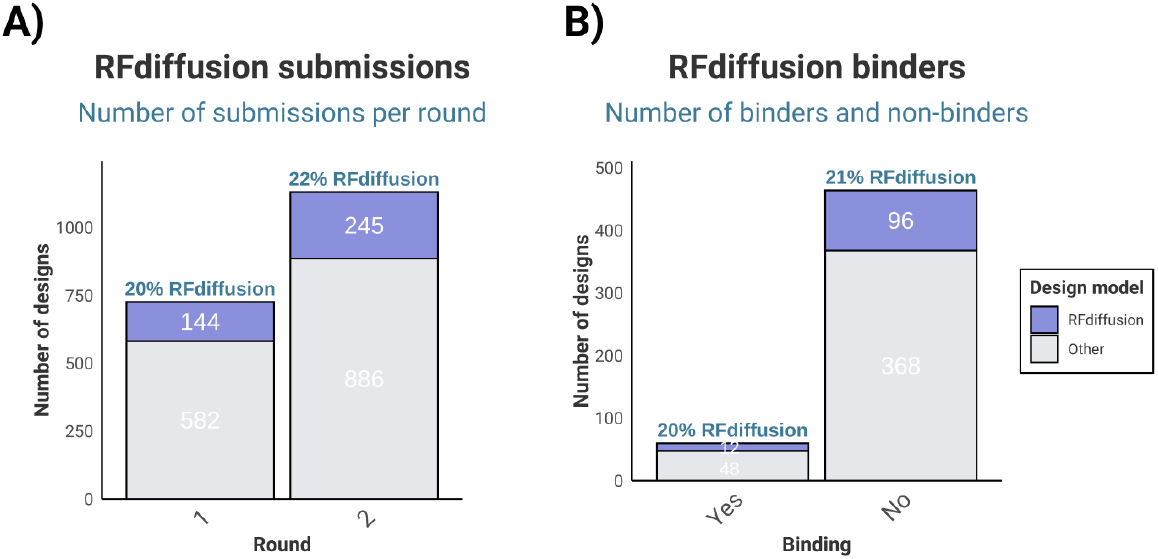
(A) Number of submissions incorporating RFdiffusion across the competition rounds, either for *de novo* or partial hallucination of existing binders, and (B) total number of RFdiffusion binders/non-binders.

A typical RFdiffusion binder design campaign begins with diffusion of binder backbones conditioned on the target structure with provided hotspot residues. This step is followed by the generation of multiple sequence candidates per binder backbone via inverse folding with ProteinMPNN [Dau+22], as RFdiffusion only generates backbone structures but not sequence, and subsequent prediction of complex structures with AlphaFold or other structure prediction models [Wat+23]. In the competition, designers sometimes incorporated additional methods to screen the binder sequences, such as PISA [KH07] and Rosetta metrics [Alf+17], while some designers expanded on the basic workflow in other ways, for instance using MD simulations to vary the target conformation prior to diffusing binders. Adrian Tripp and Sigrid Kaltenbrunner achieved third place (*K*_*D*_ 22.9 µM) in R1 with a *de novo* binder created by combining RFdiffusion backbone generation with Rosetta relaxation, LigandMPNN [Dau+23] sequence design, and sequential folding and filtering with ESMFold [Lin+23] and ColabFold [Mir+22]. Still other designers instead applied RFdiffusion’s motif scaffolding functionality to build known binding residues into new frameworks, or employed partial diffusion to develop variations of initial binders. Notable examples of binder diversification using RFdiffusion include Aurelia Bustos’s and Yoichi Kurumida’s successful binders in R2, with one of Bustos’s designs ranking fourth in affinity in R2 (*K*_*D*_ 51.6 nM).

### 3.5 ESM & Protein Language Models

Protein Language Models (pLMs) learn the probability distribution of a sequence of tokens (e.g., amino acids) after being trained in a self-supervised manner on millions of sequences. Broadly, these models can be categorized as either decoder-only pLMs (e.g., ProGen [Mad+23] or ProtGPT2 [FSH22]), typically trained with a causal modeling objective, or encoder-only pLMs (e.g., ProtT5 [Eln+20], ESM1b [Riv+21], ESM2 [Lin+23], ESM-C), typically trained with a Masked Language Modeling (MLM) objective. While the former excel at predicting the next token and are thus used for sequence generation, the latter often produce high-quality sequence representations (embeddings) [Eln+20]. An exception to this general distinction is ESM3 [Hay+25], which, despite also being trained with an MLM objective, incorporates a flexible rate that enables learning the probability distribution over all tokens and modalities, thereby allowing generation from any starting point in a single pass.

In the competition, many participants employed classical MLM-based models to design their binders (Figure 7). One team used AntiBERTy [RGS21] to propose mutations in a known scFv, then predicted and ranked these variants with DeepAB [RSG22], ultimately placing fourth and producing five (out of five) successful binders. Several contestants used ESM2 in an iterative process to maximize the log-likelihood of a starting sequence, a strategy akin to Machine Learning–Directed Evolution (MLDE). Of the top 100 *in silico* hits, 22 were generated using a modified version of this approach—three of which were successful binders, achieved by optimizing a segment of human Transforming Growth Factor alpha (TGF-α). Other teams leveraged pLMs for *de novo* sequence generation. For instance, one team fine-tuned the conditional pLM ZymCTRL [Mun+24] on a set of natural EGF-like sequences, yielding three binders [Sto+24]. Another group employed ESM3 to conditionally generate Fragment antigen-binding region (Fab)-like fragments connected by a glycine spacer, resulting in a weak binder.

**Figure 7:**
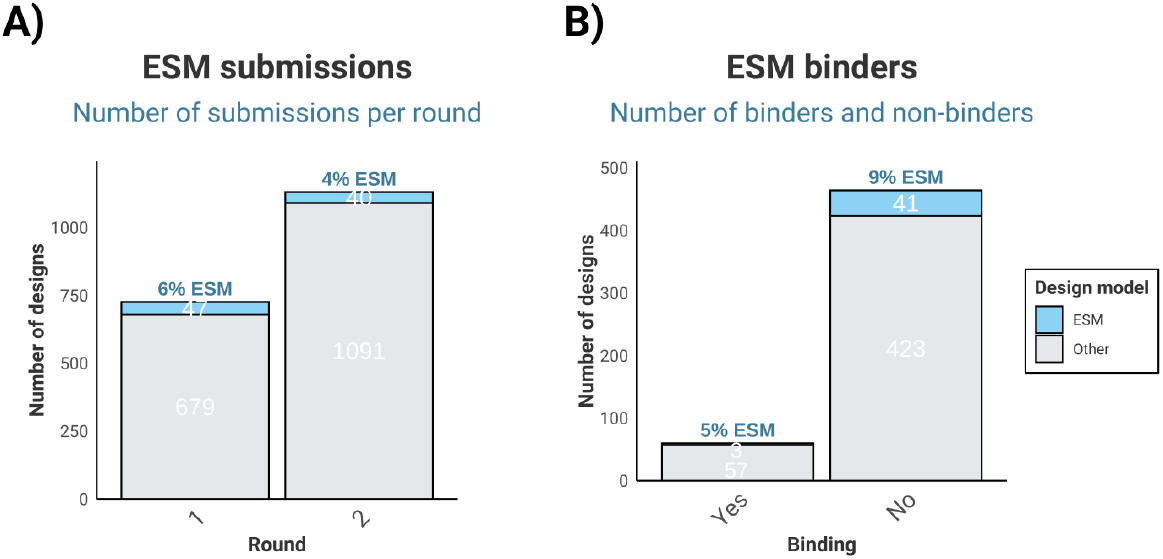
(A) Number of submissions leveraging ESM2 or 3 in their design methodology (not just for log-likelihood scoring) across the competition rounds and (B) total number of ESM binders/non-binders.

## 4 Competition Results

In this section, we provide an overview of key results from both rounds of the competition, building upon initial analyses previously presented in Adaptyv blog posts [CK24a; CK24b; CK24d]. Overall, we focus on notable improvements in design strategies, protein expression levels, and the number and quality of successful binders identified. By comparing results across rounds we illustrate how public competitions can drive methodological optimization, promote promising design approaches, uncover valuable insights, and shed light on critical gaps within the field [Bio25].

### 4.1 Round 2 yields improvements in both expressibility and binding

In Figure 8, we summarize the total number of designs selected for validation, the number of designs that expressed successfully, and the total number of validated binders across the 3 binding thresholds. We observed an improvement in the number of designs expressed, from 73% to 95%, which we attribute to knowledge gained from R1, the increased usage of ProteinMPNN with soluble weights [Gov+24] for soluble redesign, or SoluProt [Hon+21] and NetSolP [Thu+21] for predicting solubility and filtering, as well as using ESM2 log-likelihoods for scoring, biasing designs towards more naturally occurring, thus expressible, ones. We will next see if there is a direct correlation between log-likelihood scoring and expression or binding.

**Figure 8:**
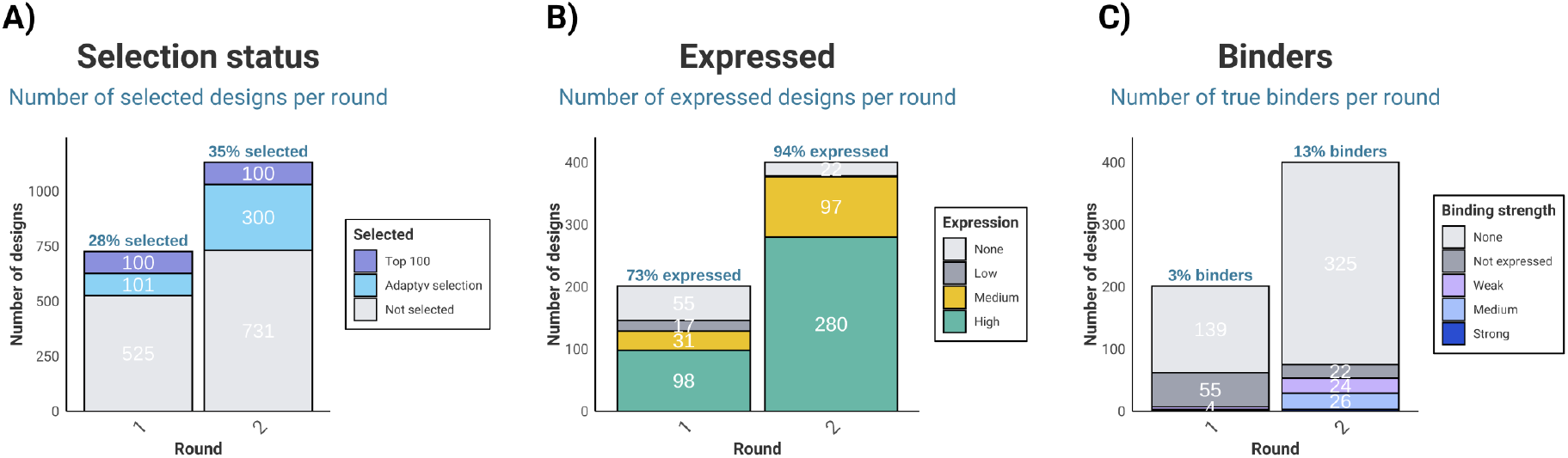
Number of (A) selected designs per round, (B) expressed designs, and (C) binders.

To assess whether any of the computational metrics used in the competition were predictive of expression, we split the binder expression data into two classes, high expression versus everything else (no expression, low, and medium expression). After this, for each of the competition’s *in silico* metrics, DE-STRESS structural metrics, and Foldseek [Kem+23] scores, we calculated the Receiver Operating Characteristic Area Under the Curve (ROC AUC) scores to understand how well these metrics could identify high expression. This analysis was performed on the structural models of the binders on their own (excluding EGFR) (Table 1).

**Table 1:**
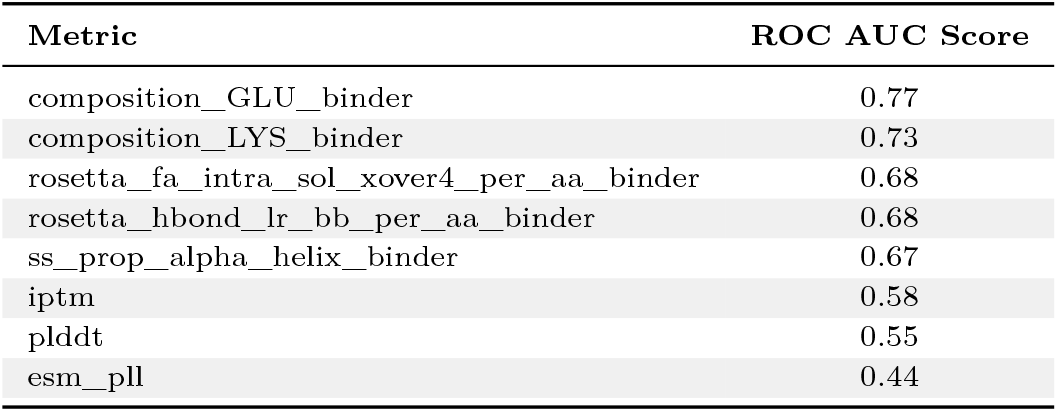
ROC AUC scores for the different metrics and expression.

The table above shows that the proportions of glutamate and lysine in the binder designs, denoted as composition_GLU_binder and `composition_LYS_binder,` achieved strong ROC AUC scores of 0.77 and 0.73, respectively. Additionally, the Rosetta all- atom scoring energies, capturing solvation `(rosetta_fa_intra_sol_xover4_per_aa)` and long-range hydrogen bonds on the backbone `(rosetta_hbond_lr_bb_per_aa)` normalised by number of amino acids, displayed relatively high ROC AUC scores of 0.68. Furthermore, the proportion of the alpha helix secondary structure within the design (ss_prop_alpha_helix_binder) had a ROC AUC score of 0.67. Conversely, the competition metrics ipTM, pLDDT and esm_pll all had relatively low ROC AUC scores of 0.58, 0.55 and 0.44, respectively, indicating they are not predictive of Adaptyv’s high-versus-other protein expression levels.

Another exciting improvement has been in the percentage of successful binders to EGFR, from 3% in the first round to 13% (Figure 8). Out of these, 2 had better binding affinities than the Cetuximab control in the nanomolar range. In total, we have amassed 60 binders, with 4 of those as strong binders (in the tens of nM range). We analyzed whether the same metrics are predictive of binding probability by repeating the DE-STRESS analysis, this time to distinguish between binding and non-binding designs. For this analysis, the DE-STRESS metrics were calculated using the AF2-M predictions for the 400 selected designs in complex with EGFR (Table 2).

**Table 2:** ROC AUC scores for the different metrics and binding.

| Metric | ROC AUC Score |
| --- | --- |
| `esm_pll` | 0.72 |
| `fident_pdb` | 0.68 |
| `alntmscore_pdb` | 0.66 |
| `pLDDT` | 0.66 |
| `evalue_pdb` | 0.64 |
| `similarity_check` | 0.64 |
| `iptm` | 0.64 |

We found that esm_pll had the highest ROC AUC score out of all the metrics, and Foldseek scores such as `fident_pdb, alntmscore_pdb, evalue_pdb,` and `similarity_check` all had ROC AUC scores of 0.68, 0.66, 0.64, and 0.64, respectively. This could be due to strategies involving optimization of existing binders having a higher probability of succeeding, and being overrepresented in the competition. Additionally, the competition metric `pLDDT` had a ROC AUC score of 0.66, while `ipTM` had a score of 0.64, indicating some degree of correlation with each of these metrics and binding probability.

Overall, these results demonstrate that while the computational metrics used for design filtering cannot accurately predict binding probabilities, the collective knowledge gained from the first round of the competition helped to improve design strategies going into R2 and increase the overall binder design success rates.

### 4.2 Shift in choice of design methods

Next, we investigate whether the design strategy changed from R1 to R2. Briefly, users were encouraged to write a short summary of their workflow, which we manually curated and categorized into five design choices:

- **Diversified binders:** These started from known binders to EGFR (such as Cetuximab, EGF, or TGF-α) and used sequence optimization tools, such as ProteinMPNN [Dau+22] or TIMED [Cas+24], to generate novel sequences that would fold similarly. Most pipelines involved self-consistency and designability filters using ESM2 log-likelihoods and AF2 refolding, scoring, and calculating distance from the original backbone (known as self-consistency).
- **Optimized binders:** Involved all active learning/Bayesian optimization loops that started from known binders, with surrogate models trained on the competition’s selection scores or known binders. These included other optimization techniques that exploit ESM2’s probabilities to increase affinity, as demonstrated before [Hie+23; Sha+24].
- **Physics-based:** Using tools such as Rosetta and MD simulations for binder diversification or even *de novo* design.
- ***de novo:*** Often diffused binders using RFdiffusion [Wat+23] or custom generative models the users developed for this application. These binders were simply conditioned on the EGFR target and started from noise.
- **Hallucination:** Included all methods using AF2 or other structure prediction networks to predict structures from a random input, further optimized via gradient descent or Markov chain Monte Carlo [Ani+21; Wic+22; Pac+24; Fra+24].

We can see that binder optimization had a 2-fold increase in the number of submissions (Figure 9 A), with a similar increase for the protein hallucination strategy. This could be attributed to the past success of tools such as BindCraft [Pac+24] that gained popularity following the first round. Among highly expressed designs, we see a major increase in *de novo* strategies, again ascribed to the increased expressibility and solubility of most submissions due to the usage of redesign tools such as ProteinMPNN with soluble weights [Gov+24]. Surprisingly, we see a sharp decrease in the prevalence of diversified designs among highly expressed sequences, with almost no changes when considering hallucinated or optimized ones - explained by the overall drop of the diversified binder strategy among all submissions. Interestingly, hallucinated designs, despite becoming more preferred in the second round, had a lower share of true binders (from 40% to 12%, Figure 9 C).

**Figure 9:**
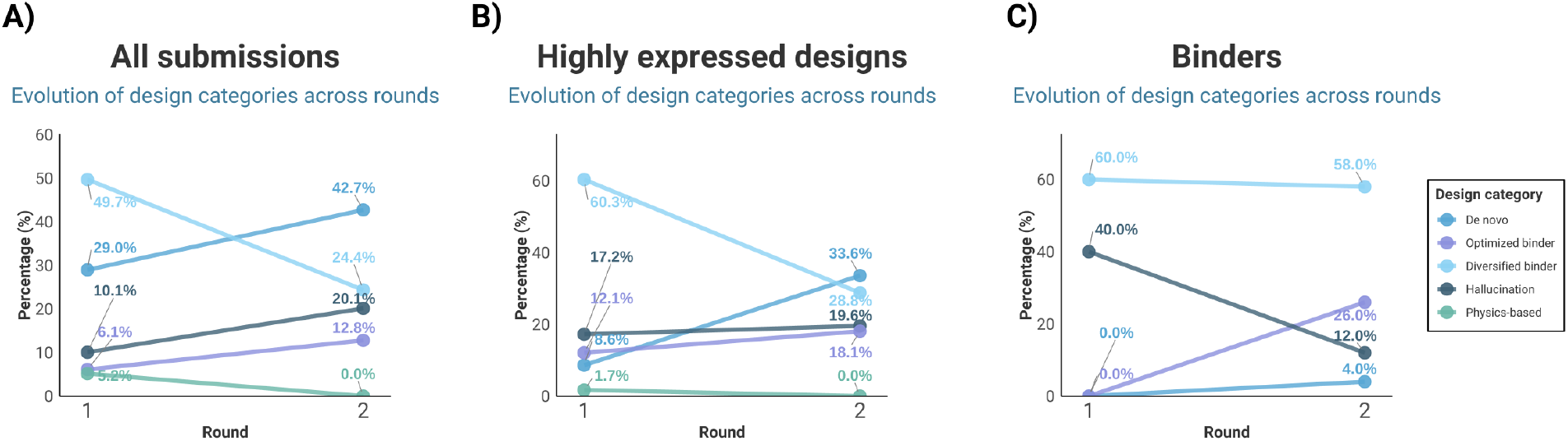
Changes in the design category (diversified binder, optimized binder, physics-based, *de novo*, hallucination) across the two competition rounds. (A) Changes across all submissions, (B) across the highly expressed designs, and (C) across experimentally validated binders.

### 4.3 Round 2 designs surpass the affinity of known EGFR binders

Figure 10 showcases all 60 true binders’ *K*_*D*_, grouped by the competition round. We see 2 binders from the first round in the tens of nM range; these were, however, disqualified afterwards, as they did not conform to our condition to be more than 10 amino acids away from a previously established EGFR binder. The next best R1 binder achieves a *K*_*D*_ on par with EGF and is a *de novo* protein designed using the BindCraft hallucination strategy [Pac+24]. Two R2 binders have *K*_*D*_ values on par or better than Cetuximab, with Cradle’s achieving an 8.2 × affinity increase compared to the control by solely modifying the framework regions [Fer+24].

**Figure 10:**
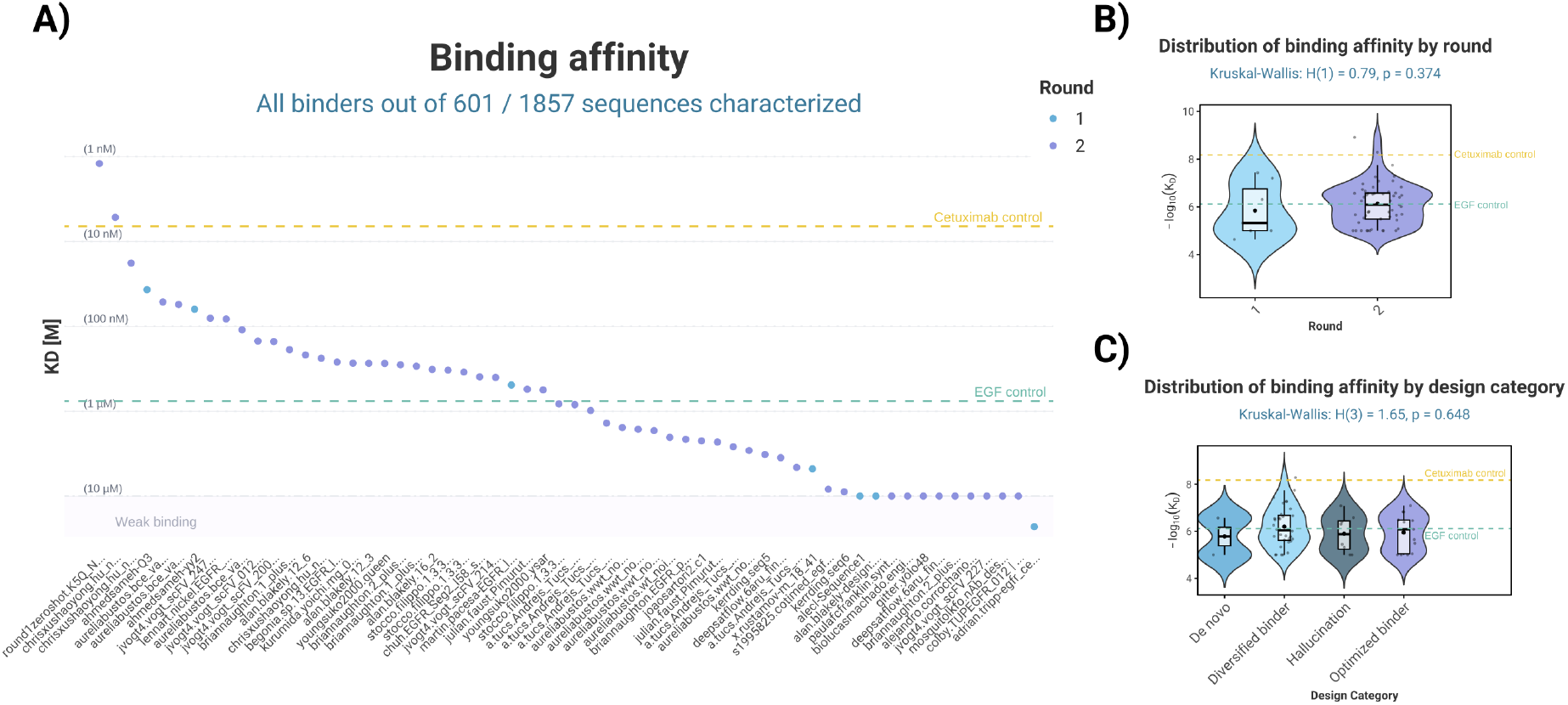
Overview of the 60 binders obtained across both competition rounds and their binding affinities. (A) Distribution of the binding affinity (averaged across 3 replicates) for each binder, (B) Distribution of binding affinities for each competition round, (C) Distribution of binding affinities across design categories.

Furthermore, we investigated whether there is an overall binding affinity improvement from one round to the next: we speculate that the prevalence of optimized binder strategies in the second round would cause a shift in the *K*_*D*_ distribution. Following a Kruskal-Wallis test, there is no statistically significant difference in the median values across the two rounds (p-value = 0.374, Figure 10 B). Similarly, there is no statistically significant difference in the median binding affinities for each design strategy (Figure 10 C). This is surprising, given the overall increase in binding success, but perhaps suggesting that current design and filtering strategies optimize for binding probability but not affinity.

Finally, we sought to better understand what influences the experimental *K*_*D*_ values of the competition binders. Figure 11 highlights the lack of statistically significant correlations between binding affinity and the metrics used for ranking designs, including the length-normalized ESM2 log-likelihood. Some unexpected trends emerge, with iPAE, ipTM, and the ESM2 likelihood showing opposite trends to what would be expected from previous studies and how the rankings were performed [Ben+23; UMG24a; CRG24]. Further analyses were performed on the ESM2 likelihood’s influence in the second round, aiming to establish a link between the inclusion of this score and the increased hit rate (from 3% to 13%).

**Figure 11:**
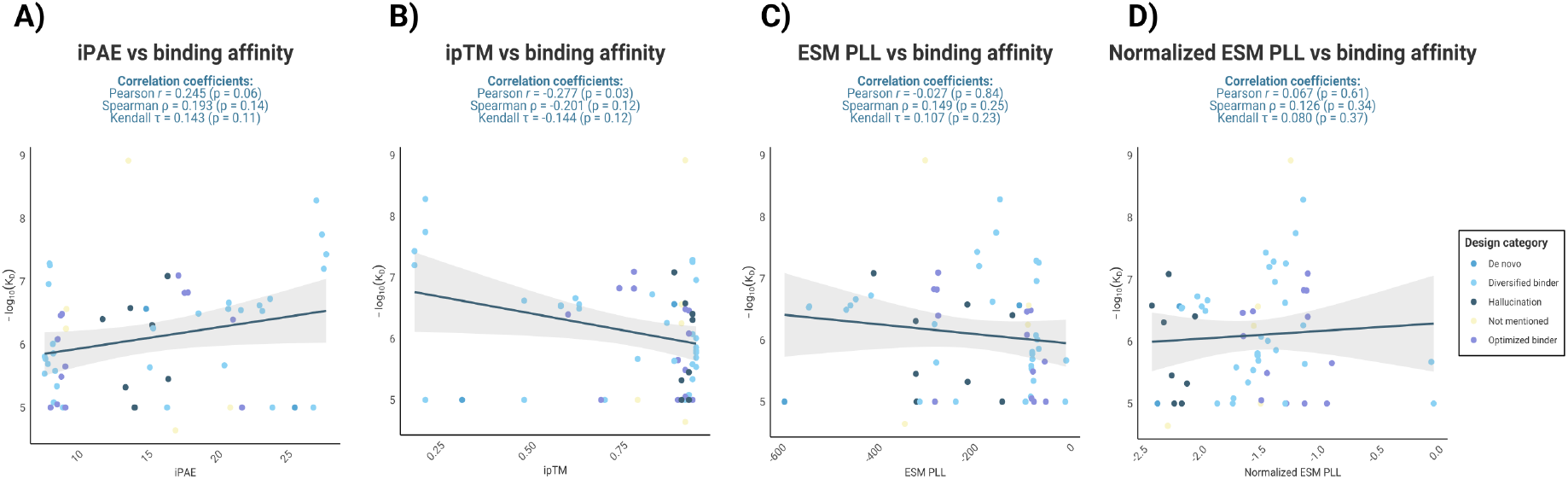
Correlation between binding affinity and the metrics used for ranking and selecting the designs. (A) iPAE correlation, (B) ipTM correlation, (C) ESM2 log-likelihood correlation, (D) ESM2 log-likelihood normalized by the sequence length.

### 4.4 Binding categories versus ESM log-probabilities

While ESM2 was primarily pretrained on non-antibody molecules, ESM3 and ESM C [Hay+25; ESM24] were released shortly after this competition and included Observed Antibody Space (OAS) sequences in their pre-training [OBD22]. Critically, in the ESM architecture, the log-probability of a sequence is derived from the sum of probabilities across each residue of a protein. Therefore, shorter proteins tend to exhibit more positive log-probabilities than longer ones, ranking more favorably.

To check whether molecule length and type were balanced prior to statistical analyses, IMGT heavy-chain renumbering was attempted on all competition sequences [Ada23; Ada23; WSK24]. Sequences were then divided into categories scFv, Nanobody, Peptide, Polypeptide, and longer proteins that failed renumbering were considered Unidentified.

In Figure 12, it can be seen that short, peptide sequences were overrepresented in the competition’s affinity characterization. Thus, we elected to balance these with 93 EGFR nanobodies characterized by Adaptyv following another public event, the 2024 Bio x ML Hackathon, to reduce bias in length-representation [Bio24; Lux]. These were designed using sequence- or structure-based methods and meet the competition’s mutation threshold at 50%–80% identity to the USPTO database. Combining the datasets, in Fig. 13 we see that unnormalized and length-normalized ESM-2 650M log-probabilities do not correlate with Adaptyv’s binder categories. There is also no relational trend for any of the unnormalized ESM model log- probabilities with *K*_*D*_. However, ESM3 and ESM C log-probabilities show strong correlation to binding categories, and even their smaller models predict better log-likelihoods—more positive—as binding strength increases.

**Figure 12:**
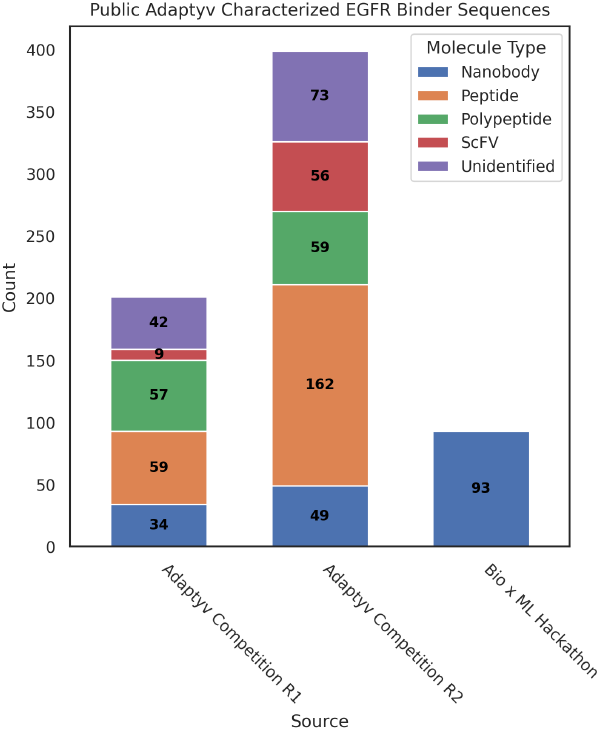
Estimated molecule counts from public, Adaptyv-characterized anti- EGFR designs.

**Figure 13:**
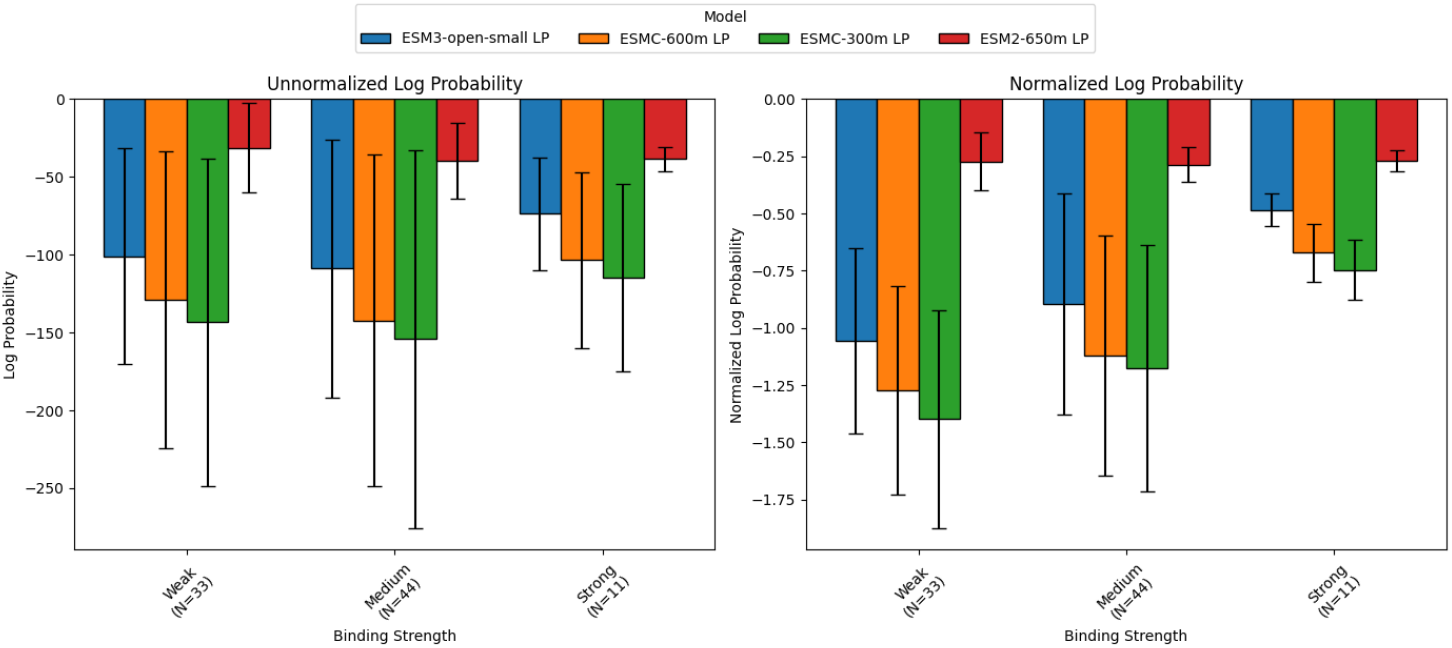
Sequence log-probabilities from ESM-2, ESM3, and ESM-C models, unnormalized and length-normalized.

### 4.5 Antibody domains require distinct structural metrics

In light of AlphaFold2’s struggles in predicting Complementarity-Determining Region 3 (CDR3) structures [Aba+23; Ken+24; CRG24; UMG24b] and its 50% or worse performance at predicting the correct epitope-paratope poses for antibodies [EG25], we elected to use the same extended EGFR dataset and molecule categories as for the ESM log-probability analysis above, which better balances the ratio of peptide and antibody-domain binders for analysis.

To check for potential biases caused by similarity to antibody-domains in pretraining datasets, we looked at the characterized Heavy Chain Complementarity-Determining Regions 3 (CDR-H3s) from the competition. We found approximately 94 unique CDR-H3s, and 16 total antibody-domain sequences displayed measurable binding (Table 3). Of the top five most common binding CDR-H3s, all led to a published therapeutic sequence with the same CDR-H3. For some of these (e.g., 4KRP), the crystal structure of the antibody-domain in complex with EGFR was known and likely included in AlphaFold2-Multimer’s training dataset.

**Table 3:**
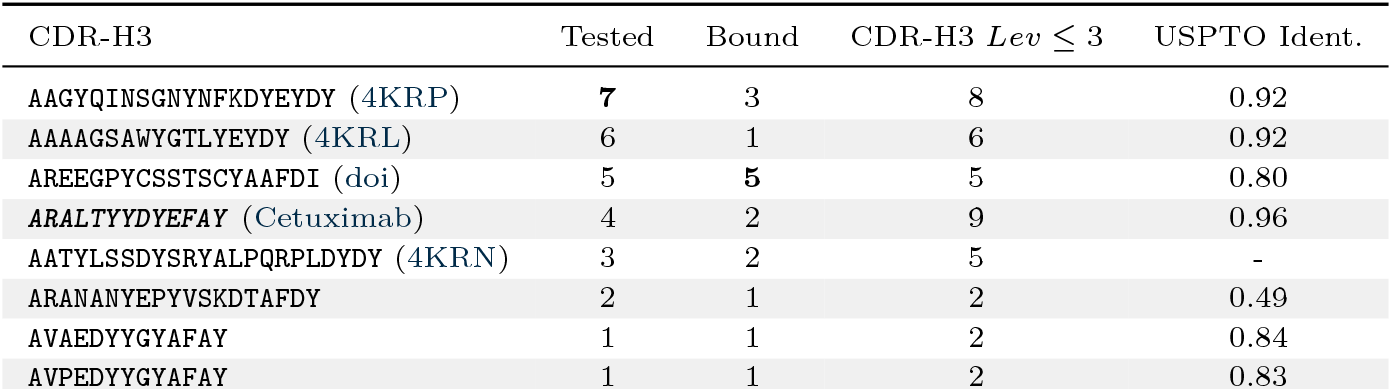
CDR-H3s of binding antibody-domain sequences from the competition; number of tested sequences with CDR-H3s within Levenshtein distance; maximum USPTO identity for full sequence.

Because of their sequence similarities to known structures or complexes, these sequences may have been posed with the correct epitope and ranked favorably by AlphaFold2 iPAE and Predicted Local Distance Difference Test (pLDDT) metrics. Additionally, these known structures would have allowed participants to take an existing binder and fold a new sequence to the same structure, making them attractive choices for structure-informed design submissions.

Some binding CDR-H3s, such as ARANANYEPYVSKDTAFDY, had no readily apparent parent molecule, and were fairly unique in terms of overall sequence identity (Table 3). However, while they technically exhibited measurable binding, they were Weak binders, close to the limit of detectable binding and likely not suitable for further development.

Next, in order to correlate predicted structural metrics with *K*_*D*_, we evaluated the results from AF2-M, mpDockQ [Bry+22], and the PI-score pipeline [Mal] using AlphaPulldown [Yu+23].

Table 4 compares the ANOVA F-stats for binders and non-binders using metrics derived from each of the models and ESM-2/C (see A.6 for metric descriptions).

**Table 4:** ANOVA F-stats for Binders and Non-Binders. Stats with significant p-values approximately ≤ 0.0003 bolded; top three double-digit F-stats italicized for each column. *Peptide* and *Ab-Domain* are competition sequences, *Ab-Domain Extended* includes the Bio x ML Hackathon EGFR nanobodies, and *All + Unidentified* includes all previous columns plus longer and Unidentified competition proteins. P-values are unadjusted for multiple comparisons.

|  | All + Unidentified | Ab-Domain Extended | Peptide | Ab-Domain |
| --- | --- | --- | --- | --- |
| ESM-2 logP scaled | **29.919** p=6.4e-08 | 0.047 p=0.83 | **30.360** p=1.0e-07 | 0.085 p=0.77 |
| ESM C logP scaled | **27.767** p=1.9e-07 | 0.246 p=0.62 | **8.690** p=3.6e-03 | 0.052 p=0.82 |
| ipTM-pTM | 0.038 p=0.85 | 2.362 p=0.13 | **13.235** p=3.4e-04 | 3.720 p=5.7e-02 |
| ipTM | 0.038 p=0.85 | 2.474 p=0.12 | **12.679** p=4.6e-04 | 3.884 p=5.2e-02 |
| pDockQ/mpDockQ | **11.089** p=9.2e-04 | **9.862** p=1.9e-03 | **19.443** p=1.7e-05 | **18.532** p=3.9e-05 |
| avg iPAE | 0.200 p=0.66 | 2.885 p=9.1e-02 | **12.795** p=4.3e-04 | 4.651 p=3.3e-02 |
| avg ipLDDT | 4.235 p=0.040 | **7.518** p=6.6e-03 | **8.812** p=3.3e-03 | **11.601** p=9.5e-04 |
| binding energy | **23.160** p=1.9e-06 | 6.423 p=0.012 | **13.045** p=3.8e-04 | 3.035 p=8.5e-02 |
| num intf AAs | 0.670 p=0.41 | 0.200 p=0.66 | 2.900 p=9.0e-02 | 0.822 p=0.37 |
| polar | **7.514** p=6.3e-03 | 0.104 p=0.75 | 0.115 p=0.74 | 0.173 p=0.68 |
| hydrophobic | **6.877** p=8.9e-03 | 1.319 p=0.25 | 0.463 p=0.50 | 3.812 p=5.4e-02 |
| charged | 2.495 p=0.12 | **11.902** p=6.7e-04 | 0.028 p=0.87 | **17.382** p=6.6e-05 |
| contact pairs | 0.149 p=0.70 | 1.802 p=0.18 | 1.712 p=0.19 | 1.198 p=0.28 |
| sc | 1.152 p=0.28 | 4.771 p=0.030 | **15.172** p=1.3e-04 | 5.887 p=1.7e-02 |
| hb | 4.871 p=2.8e-02 | 0.255 p=0.61 | **8.205** p=4.6e-03 | 1.334 p=0.25 |
| sb | 0.882 p=0.35 | 1.808 p=0.18 | **13.756** p=2.7e-04 | 5.296 p=2.4e-02 |
| int solv energy | 0.566 p=0.45 | 5.291 p=2.2e-02 | **7.902** p=5.4e-03 | 1.136 p=0.29 |
| int area | **6.945** p=8.6e-03 | **10.970** p=1.1e-03 | **14.289** p=2.1e-04 | **8.203** p=5.1e-03 |
| pi score | 1.331 p=0.25 | 1.559 p=0.21 | **18.649** p=2.4e-05 | 3.044 p=8.4e-02 |

We find that `pDockQ/mpDockQ` scores and `int area` seem to be relevant for all binding molecules, with a larger interface area corresponding to binders. For *Peptides*, length normalized ESM log-probabilities, `sc,` and `pi` score also have relatively high F-stats, along with `avg iPAE` and `iPTM-pTM.` However, these metrics and others relevant to *Peptides* do not seem to be relevant for antibody-domain binders. Metrics like `charged` and `avg ipLDDT` seem to correlate with binding for the *Ab-Domain* competition sequences, but also have less significant F-stats and p-values when considering the larger dataset of *Ab-Domain Extended*. In that group, the relatively significant F-stats (but greater p-values) for `charged` and `int area` reinforce the relevance of those metrics for all Adaptyv-characterized EGFR antibody-domain molecules.

Notably, when evaluating all the molecule types together, the F-stats can paint a misleading picture of which metrics might inform binder design. For instance, `binding energy` and ESM logPs have high F-stats in the *All + Unidentified* column, but these metrics only seem to distinguish binders from non-binders for the *Peptide* group. Similarly, `pDockQ/mpDockQ` has an F-stat of about 11.1 for *All + Unidentified*, but it has a more significant p-value and F-stat of 19.4 and 18.5 for `Peptide` and `Ab-Domain,` respectively, and a considerably insignificant p-value and F-stat of 9.9 in `Ab-Domain Extended.` This highlights the need for sufficient sample sizes, along with domain- and molecule-specific metrics when performing computational protein design.

## 5 Discussion & Perspectives

The Adaptyv EGFR binder competition provided a unique and valuable opportunity for the computational protein design community to benchmark diverse methodologies in a standardized, unbiased, and collaborative setting. By offering an open, standardized experimental platform, the competition enabled researchers from various institutions and backgrounds to test and compare methods without the typical constraints and delays associated with traditional publishing. Such initiatives are especially important in protein design, where methodological innovations rapidly evolve, and experimental validation remains both critical and resource-intensive [Arm+24].

Additionally, the availability of open-source computational pipelines (e.g., Brian Naughton’s biomodals) and cloud computing credits provided by Adaptyv’s partner modal.com greatly broadened participation. Together, these factors fostered transparency, rapid innovation, and strong community engagement, highlighting the value of collaborative efforts in driving scientific progress.

### 5.1 Limitations of current computational metrics

One primary insight from the competition is that, while significant advances have been achieved in computational protein binder design, the challenge remains far from solved. Deep learning-based design methods, such as RFdiffusion or BindCraft, while achieving high levels of experimental success, can still produce many false positive designs with high *in silico* scores. This outlines clear limitations in the currently used computational metrics and methods used for design filtering, particularly regarding the prediction of binding probability or affinity. While numerous affinity prediction models currently exist, ranging from physics-based and simple ML models [Xue+16] to complex DL ones [MBB24] or likelihoods from pre-trained pLMs [Lin+23], none of them had been confirmed to strongly generalize at the time of the competition [CRG24]. The ranking metrics used, such as AF2 iPAE and ipTM scores, provided valuable preliminary screens but were insufficiently predictive of true experimental binding probability or affinity (see section 4.5).

This suggests these metrics are perhaps not best suited as final predictors for choosing designs for experimental characterization, as one could potentially filter out a large amount of viable binders. This can likely be attributed to the fundamental nature of these metrics, derived from structural confidence scores, which do not fully represent all biochemical and biophysical aspects one would expect of a good binder. Additionally, none of the deep learning-based methods were trained on datasets informed by binding affinity, rather relying on networks trained on co-evolutionary information and the presence or absence of protein complex structures of varying affinities in the PDB.

The models and datasets also suffer from a lack of uniformity in training data due to variability in experimental conditions and assays, such as yeast display, Chromatin Immunoprecipitation Sequencing (ChIP-seq), BLI, and Surface Plasmon Resonance (SPR), which can contribute to their poor performance. Addressing this gap requires standardized deposition and thorough curation of affinity measurement datasets. The datasets generated by this and similar competitions could serve as invaluable resources for training and benchmarking such predictive models, by providing a large and standardized resource with a variety of different targets.

### 5.2 Challenges in defining binding hits

Defining a true binding hit remains a nuanced challenge. At sufficiently high concentrations, nearly any pair of molecules can interact, making clear, standardized criteria based on robust experimental benchmarks essential. Discrepancies between studies highlight the problem: RFdiffusion classified any binder reaching 50% of the positive control signal on BLI as a hit, complicating evaluation when no strong positive control exists. In contrast, BindCraft counted hits as any binder with a clear SPR signal at 10 *µ*M. These critical differences in defining success rates underscore the need for a community consensus on experimental validation thresholds to ensure consistency and comparability across computational methods.

### 5.3 The need for standardized benchmark targets

Another significant takeaway is the need for standardized benchmark targets within the computational protein design community. Certain targets, such as Programmed Death-Ligand 1 (PD-L1), have begun to emerge as common benchmarks [Wat+23; Pac+24; Zam+24], due to their ideal and well-characterized binding site, enabling high success rates for binder design. For example, the BindCraft paper reports a 66% *in silico* success rate for PD-L1, compared to less than 10% for other targets. However, most commonly used targets are of human origin, which poses challenges for efficient production, stability, structural characterization, presence of post-translational modifications, and flexibility. These obstacles make it difficult to systematically assess and compare different computational methods. Therefore, selecting a set of standard benchmarking targets that addresses the challenges of *in silico* design, experimental validation, and structural characterization would be highly beneficial for driving the field forward.

To address this need, we propose the Bench-tested Binder Benchmark (BenchBB), a curated set of standardized protein-binding targets designed to enable rigorous, consistent, and practical evaluation of computational binder design methods.

### 5.4 BenchBB: a standardized benchmark for protein binder design

Outside of protein design, the ML field has iterated through various benchmarks as models progressed and saturated (e.g., GLUE [Wan+18], SuperGLUE [Wan+19], MMLU [Hen+20]), developed specialized variations for specific purposes (e.g., imagenet1k [Den+09] as an economical computer vision benchmark), and addressed shortcomings in benchmark design (e.g., SWE+ [Ale+24] addressing limitations of the SWE benchmark [Jim+23]). Similarly, protein design models could face limitations, such as poor generalization to unseen Protein-Protein Interactions (PPIs), failure to identify highest-fitness variants, or exploitation of assay-specific idiosyncrasies (reward hacking), thereby hindering broader applicability.

We consider BenchBB as a practical starting point in the development of standardized benchmarks that consider these challenges and expect that it will be improved upon as the field develops. BenchBB includes the following targets, selected in an attempt to balance experimental accessibility, structural characterization potential, and diversity in binder modalities ^2^ :

- **PD-L1:** De facto binder design benchmark target [Wat+23; Pac+24; Zam+24]. Can be produced via secreted human cell expression or through refolding from *E. coli*.
- **EGFR:** Ectodomain easily produced in human cells; directly comparable to results from the Adaptyv EGFR binder competition.
- **IL7Ra:** Ectodomain easily produced in human cells; benchmarked in multiple prior studies [Wat+23; Cao+22; Zam+24].
- **BHRF1:** Introduced as a target in the AlphaProteo paper [Zam+24]; easily expressed in *E. coli* and commercially available with many antibody controls.
- **SpCas9:** CRISPR nuclease that enables easy structural characterization via cryoEM; stable, easily expressed in *E. coli*, and with multiple known binding sites and conformations.
- **BBF-14:** A *de novo* designed protein [Gov+24] that can assess generalization beyond natural interfaces; bench- marked in BindCraft, features two characterized binding interfaces, and is easily produced in *E. coli* [Pac+24].
- **MBP:** Highly soluble, easily expressed in *E. coli*, and widely used as a fusion tag to enhance solubility. Features a well-characterized active site allowing straightforward binder screening via elution from amylose resin.

Like most benchmarks, BenchBB represents a minimum performance threshold: methods that fail to yield measurable-affinity binders after testing a practical number of variants are unlikely to outperform the current SOTA. Conversely, methods demonstrating strong results on BenchBB should still avoid exaggerated claims of broad generalization or assertions that they have “solved” protein binder design.

Choosing the optimal design strategy, whether *de novo* generation or optimization of existing binders, remains highly context-dependent and requires significant domain knowledge. For example, despite RFdiffusion’s popularity in R2 for *de novo* design, top-performing designs typically optimized or diversified existing binders (see sections 3.4 and 4.2). Selecting appropriate starting sequences, targeting specific binding sites, and interpreting scoring metrics correctly demands careful consideration of both biophysical constraints and model biases. As shown in section 4.5, different domains and molecule types may necessitate distinct interpretations (e.g., high `binding energy` thresholds differing between peptides and larger molecules).

Finally, domain context also influences the suitability of a design for a potential application. Peptides and miniproteins, while valuable in basic research, still face challenges in therapeutic translation due to unresolved concerns such as immunogenicity. Conversely, nanobodies and antibodies with proven clinical potential are computationally demanding to design, due to limited co-evolutionary signals and reliance on loop-dominated interactions, which are poorly captured by existing deep learning models such as AF2.

## 6 Conclusion

Looking forward, competitions like the Adaptyv EGFR binder challenge have the potential to significantly advance the computational protein design field by providing expansive, validated datasets, encouraging standardization of methods and benchmarks, prospectively evaluating predictions, and promoting collaborative scientific inquiry. The introduction of standardized benchmarks such as BenchBB will further strengthen this impact, allowing consistent, rigorous evaluation and direct comparisons across emerging computational methods. Beyond binding and affinity, future competitions could incorporate additional criteria reflecting practical therapeutic or industrial considerations, such as biological signaling outcomes, immunogenicity, solubility, and environmental stability. Such multi-objective optimization tasks would more accurately reflect real-world protein design challenges, thereby driving the development of more robust and practical computational design methodologies. Whatever variations and changes are introduced in future competition formats, however, it remains crucial that competitions, associated pipelines (both models and code), and resulting datasets remain as open and accessible as possible. We expect this combination of frictionless reproducibility [Don23], standardized benchmarks like BenchBB, and increasingly meaningful competition targets will serve as the key drivers of continued innovation in computational protein design.

## 7 Code and Data Availability

Software used in the Adaptyv competition and data packages are available from

- https://github.com/adaptyvbio/egfr_competition_1
- https://github.com/adaptyvbio/egfr_competition_2
- https://github.com/adaptyvbio/competition_metrics
- https://github.com/adaptyvbio/egfr2024_post_competition

Participants’ protein design method descriptions and visualizations are also available at https://foundry.adaptyvbio.com/competition.

## 8 Acknowledgments

M.P. was supported by the Peter und Traudl Engelhorn Stiftung. B.E.C. was supported by the Swiss National Science Foundation, the NCCR in Chemical Biology, the NCCR in Molecular Systems Engineering. L.N. was supported by the Novartis Foundation for Medical-Biological Research. L.V.C. is supported by the UK Research and Innovation (UKRI) Centre for Doctoral Training in Biomedical AI at the University of Edinburgh (EP/S02431X/1). T.K. was funded by a University of Edinburgh School of Biological Sciences PhD Scholarship. K.S. was funded by the Royal Society’s University Research Fellowship. S.M.U. is supported by the UKRI Biotechnology and Biological Sciences Research Council (BBSRC) grant number BB/T00875X/1 under EASTBIO DTP). M.J.S. and C.W.W. were supported by UK Research and Innovation (UKRI) Biotechnology and Biological Sciences Research Council Strategic Longer and Larger award (BB/X003027/1). C.N.C. and F.P. were supported by the Air Force Office of Scientific Research under award FA9550-20-1-0351; C.N.C. was also supported by the San Diego Fellowship. A.G. was supported by the National Science Foundation award 2226451, computing resources of the University of Wisconsin-Madison Center for High Throughput Computing [Cen06], and generous support from Jeanne M. Rowe. L.H. was supported by the Unidel Distinguished Graduate Scholars Fellowship and NSF NAIRR Pilot 240064.

## Acronyms

AF2: AlphaFold2. 2, 5, 9, 10, 13, 15
AF2-M: AlphaFold2 Multimer. 3, 5, 12
AFDB: AlphaFold Protein Structure Database. 5
BenchBB: Bench-tested Binder Benchmark. 14, 15
BLI: Bio-Layer Interferometry. 1–3, 5, 6, 14
CDR3: Complementarity-Determining Region 3. 12
CDR-H3s: Heavy Chain Complementarity-Determining Regions 3. 12
CDRs: Complementarity-Determining Regions. 5
ChIP-seq: Chromatin Immunoprecipitation Sequencing. 14
CNNs: Convolutional Neural Networks. 6
DL: Deep Learning. 2, 13
EGF: Epidermal Growth Factor. 6, 7, 9, 10, 26, 27
EGFR: Epidermal Growth Factor Receptor. 1–3, 5, 6, 8–13, 15, 25–27
Fab: Fragment antigen-binding region. 7
IGF1R: Insulin-like Growth Factor 1 Receptor. 26
iPAE: Interface Predicted Aligned Error. 3, 6, 10–13
ipTM: Interface Predicted Template Modeling Score. 3, 5, 6, 10, 11, 13
MD: Molecular Dynamics. 6, 7, 10
ML: Machine Learning. 1, 2, 5, 13, 14
MLDE: Machine Learning–Directed Evolution. 7
MLM: Masked Language Modeling. 7
MSA: Multiple Sequence Alignment. 3
OAS: Observed Antibody Space. 11
PDB: Protein Data Bank. 5, 14
PD-L1: Programmed Death-Ligand 1. 14
pLDDT: Predicted Local Distance Difference Test. 12
pLM: Protein Language Model. 5, 7
pLMs: Protein Language Models. 7, 13
PPI: Protein-Protein Interaction. 14
R1: Round 1 of the Adaptyv Bio EGFR Binder Design Competition. 3–7, 9, 10, 26
R2: Round 2 of the Adaptyv Bio EGFR Binder Design Competition. 3–7, 9, 10, 15, 26
ROC AUC: Receiver Operating Characteristic Area Under the Curve. 8, 9
scFv: Single-chain Variable Fragment. 5, 7, 11
SOTA: State-Of-The-Art. 6, 15
SPR: Surface Plasmon Resonance. 14
TGF-α: Transforming Growth Factor alpha. 7, 9, 27

## A Supplementary Results

We include additional analyses that did not fit the flow of the main body to further contextualize thos results.

### A.1 Score distributions shifted across the 2 competition rounds

We investigate whether, across all submissions, there was an improvement in the overall scores used for ranking designs (iPAE in the first round, with the log-likelihood of ESM2 and ipTM added in the second round) or other sequence properties. There is a distribution shift for the unnormalized ESM2 score (biased against long sequences, Figure 14 A), caused by the prevalence of smaller designs (peak at the 50 residue length, Figure 14 C) in the second round. We see, however, a positive shift in the ESM PLL normalized by sequence lengths for the first round, where redesigning existing binders was the widespread choice (thus a higher PLL due to sequence naturalness/in-distribution bias) compared to the *de novo* or hallucination strategies. Another slight shift can be observed in the number of interface residues, with the second round seeing an increase in hallucination strategies that incorporated this metric in the loss function [Pac+24]. Lastly, we see some ipTM peaks at 1.0 in the second round, explained by the fact that this became a selection metric compared to the first round, and overall lower iPAE values in the second round. This indicates that most users improved their design strategies to conform to the new selection methods, yet we cannot be certain these changes caused the increased number of expressed and binding designs seen in the second round.

**Figure 14:**
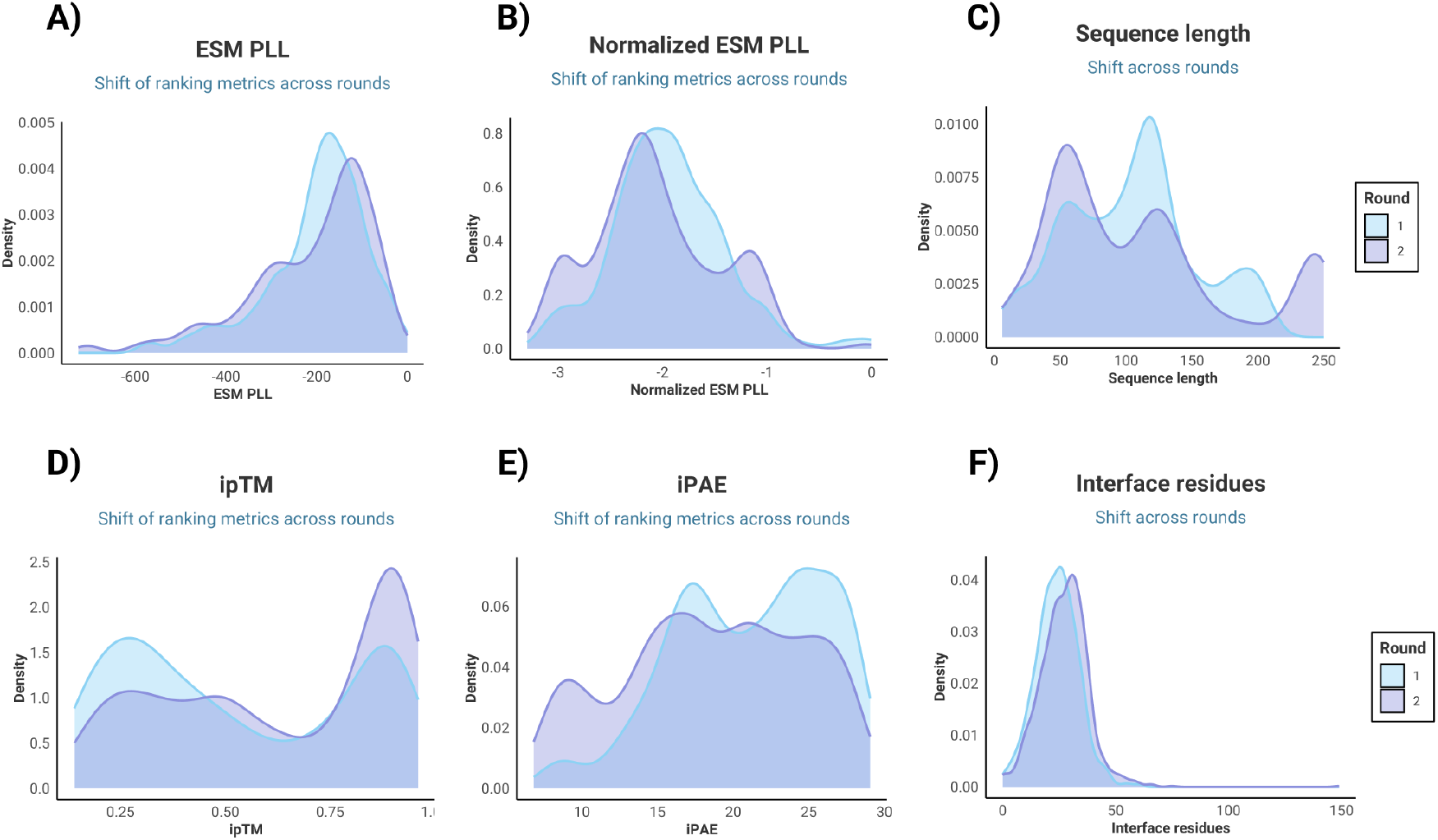
Distribution (KDE) shift of several computational scores used for design ranking and selection, for all submissions. (A) For the ESM2 pseudolikelihood, unnormalized, (B) ESM2 PLL normalized by the sequence length, (C) sequence length distribution, (D) Interface predicted template modeling (ipTM) score, (E) Interface predicted alignment error (iPAE), (F) Number of interface residues.

### A.2 Amino acid compositions do not differ drastically among *de novo* or existing binders compared to non-binders

We assessed whether the binder amino acid compositions (both at the whole sequence and at the interface level) change compared to non-binders (Figure 15 A). We see an enrichment of acidic amino acids (glutamic acid, 14.5% for *de novo* interfaces, 10.6% for existing binder interfaces and 10.2% for non-binders) for *de novo* binders at the interface and whole sequence levels (with 21.4% in *de novo* and only 16% in non-binders, Figure 15 B). Again, this can be ascribed to redesign methods that aim to increase the solubility of *de novo* designs. Tyrosine is more enriched in the existing binder interfaces (10%), compared to *de novo* binders, while glycine appears more frequently at the whole sequence level. We do not see an enrichment of hydrophobic residues at the interface level compared to the whole sequence, as one might expect for most (*de novo*) binders (Figure 15 C), and as exemplified before for target structure-conditioned designs [Cao+22].

**Figure 15:**
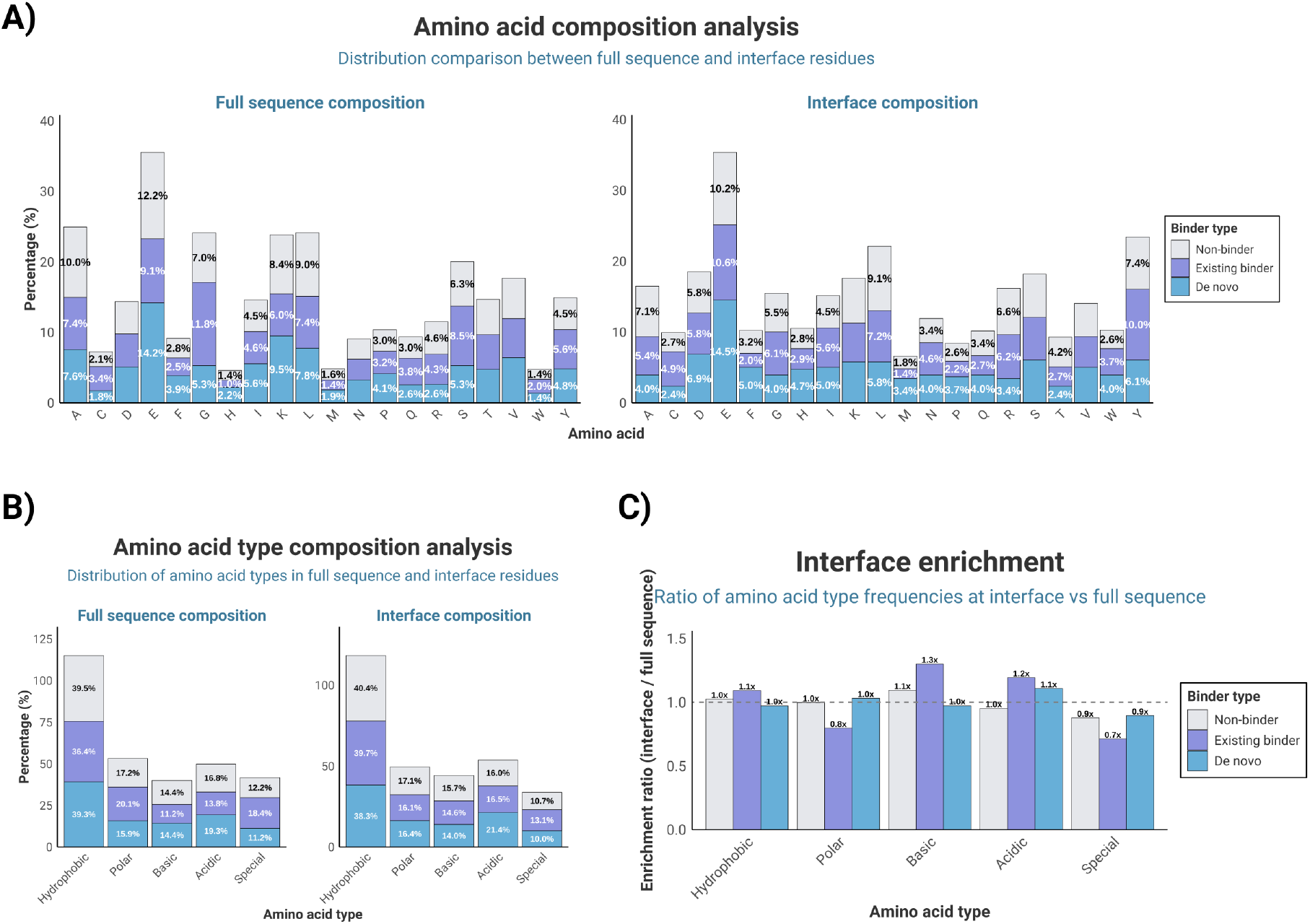
Amino acid composition analysis for the validated binders, split by *de novo* binders, existing binders (optimized, diversified), and non-binders. (A) Amino acid composition for both the full design and interface region, (B) Composition by amino acid type, (C) Enrichment ratios (at the interface versus full sequence) by amino acid type.

**Figure 16:**
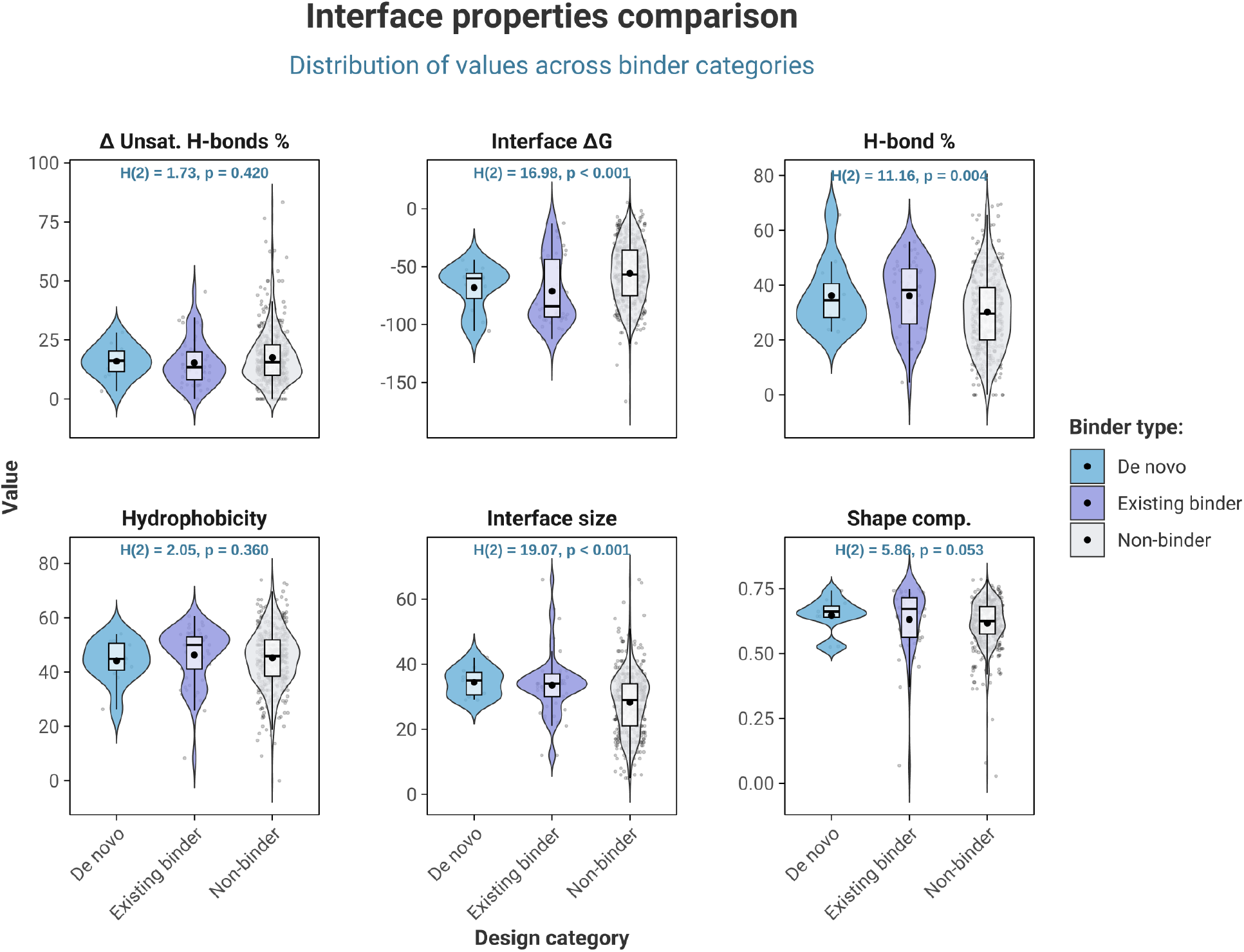
Distribution of several interface physicochemical properties by binder type (*de novo*, from existing binder, or non-binder). Properties include: *Δ* unsaturated H-bonds %, interface *Δ*G, percentage of H-bonds, hydrophobicity, number of interface contacts, and shape complementarity score.

### A.3 inding interfaces have more hydrogen bonds, better *ΔG*, and an increased number of interface contacts

Finally, we compared the interface physicochemical properties for designs starting from known binders versus *de novo* and non-binders. The main differences are across the interface *ΔG* (different median values between binders and non-binders, with a p-value *<* 0.001 following a non-parametric Kruskal-Wallis test), the number of interface contacts (interface sizes, p-value *<* 0.001), and percentage of hydrogen bonds at the interface (p-value = 0.004). *De novo* binders demonstrate more favourable binding free energies compared to existing binders, which might be caused by the rigid *ΔG* filtering criteria for these designs. Binding interfaces also display an increased shape complementarity, especially the *de novo* designs. There are no statistically significant differences in the hydrophobicity median values.

Overall, this indicates that valid binding interfaces are characterized by favourable free energies, an increased number of contacts, and the formation of H-bond networks. These could be emphasized in any binder design protocol, with many having implemented binding energy or shape complementarity filtering thresholds [Pac+24].

#### A.3.1 Cross-Correlation Analysis

First, we analyzed the relationships using Pearson correlation between the features to identify variables that potentially explain similar underlying properties.

As shown in Figure 17, `composition_LYS_binder, composition_GLU_binder, ss_prop_alpha_helix_binder, and rosetta_hbond_lr_bb_per_aa_binder` exhibit correlations with one another and negative correla- tions with similarity_check, which range from R = −0.43 (`composition_LYS_binder`) to R = −0.69 (rosetta_hbond_lr_bb_per_aa_binder). This suggests that designs that had better `rosetta_hbond_lr_bb_per_aa_binder` were also closer in sequence similarity with known binders, exhibiting similar hydrogen bonds.

**Figure 17:**
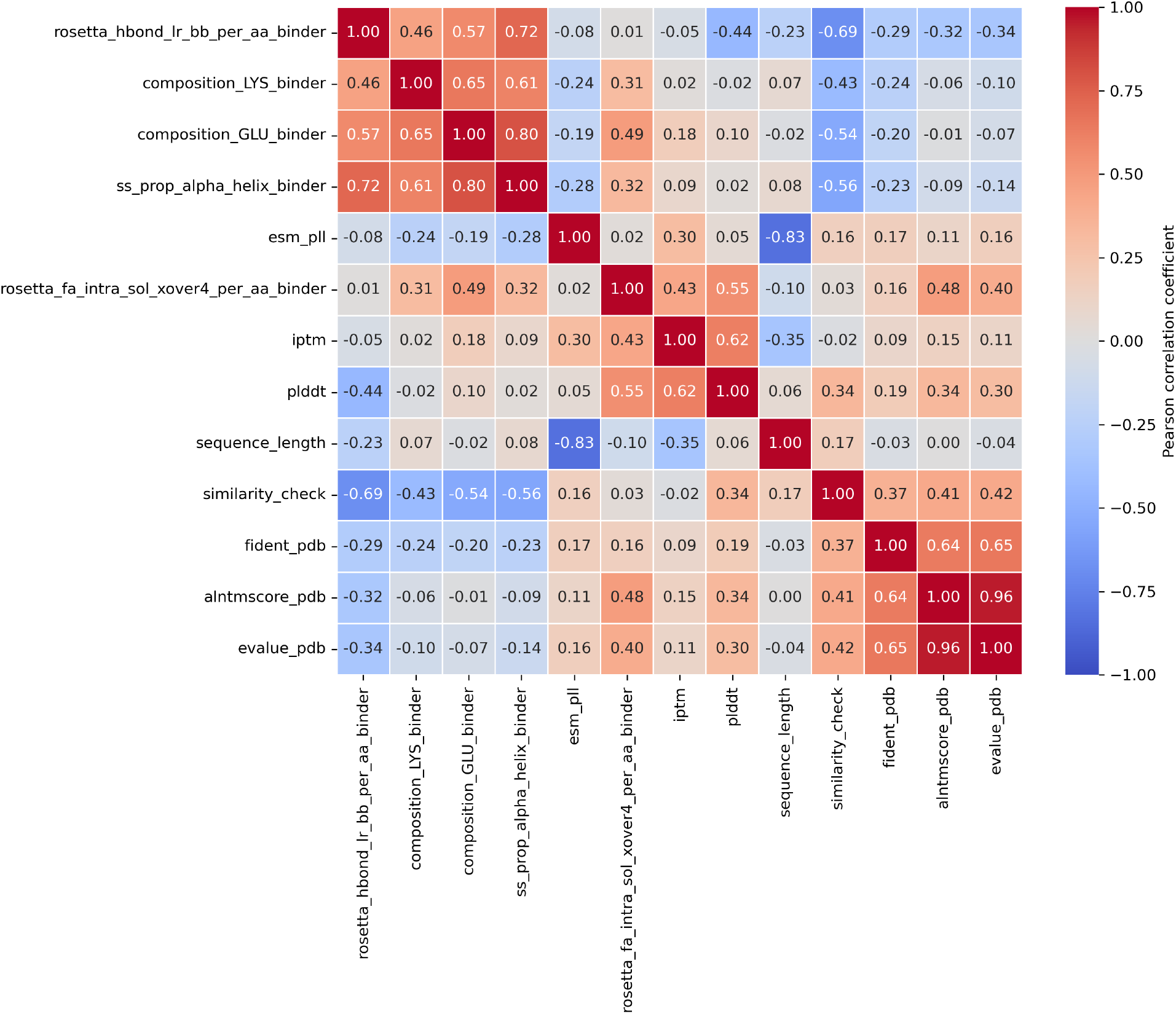
Cross-correlation matrix of the main binding and expression predictors. The heatmap illustrates the correlations between features calculated from binders’ sequences and structures. The color gradient represents the strength and direction of these relationships, with red indicating positive correlations and blue indicating negative correlations.

`fident_pdb, alntmscore_pdb`, and `evalue_pdb` show weak correlations with `pLDDT` and `similarity_check` (R = 0.37 to 0.42), and are strongly correlated between themselves (from R = 0.64 to 0.96). This points to a limited but existing redundancy in what they could be explaining about the data.

Another important predictor, AlphaFold’s `pLDDT,` correlates with rosetta_hbond_lr_bb_per_aa_binder (R = −0.44), `rosetta_fa_intra_sol_xover4_per_aa_binder` (R = 0.55), and iptm (R = 0.62). Additionally, `sequence_length` is a strong predictor of `esm_pll,` as expected, given that the latter is not normalized by length.

Coupled with the results in Tables 1 and 2, these findings suggest that while a combination of physico-chemical and deep learning-based scores could be used to filter sequences, there is also a high degree of redundancy among the descrip- tors—potentially due to confounding factors such as sequence similarity and biases in the design choices. Meaning that this can limit the generalizability of certain predictors, particularly for features highly correlated with `similarity_check.`

### A.4 Correlation between DNA features and expressibility

Using the features in Fig. 18, a binary classifier was created for High, Medium, and Low expression as the Positive class, and No Expression as the Negative class. The features were obtained from DNAChisel and primarily consist of string-based and arithmetic features. We were able to obtain a classifier with an *F* 1 of 0.93, and an accuracy of 90%.

**Figure 18:**
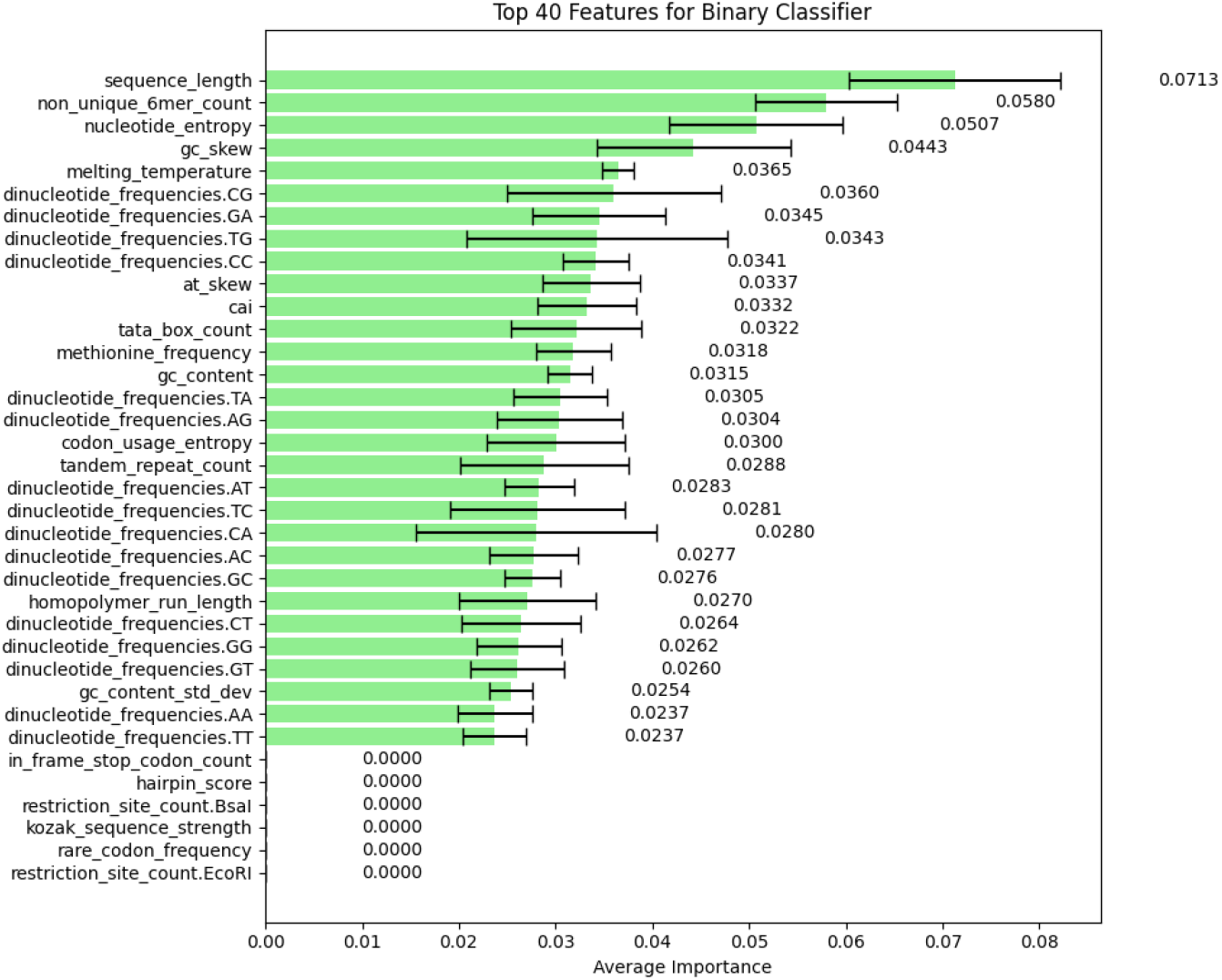
Importance of each DNA feature for expression, given by gain in model accuracy.

We found that sequence length is the most significant feature for expression, contributing a 7.13% gain in model accuracy. Non-unique 6-mer count, a representation of repeats was the next most significant feature, followed by features like entropy, skew in GC content, and melting temperature.

Due to the similar accuracy of predicting expression using either amino acid sequences or DNA sequences, we conclude that the DNA synthesis process produced no confounding expression issues for the submitted amino acid designs. Thus, expression was primarily related to the submitted sequence and was in the hands of the designer.

### A.5 EGFR Contact Analysis

Based on the AlphaFold predictions, we assessed which EGFR residues were most often targeted by designers and whether any specific target contacts were more prevalent among successful binders. For the purpose of this analysis, we defined the contact frequency for a given EGFR residue as the fraction of a set of designs in which that EGFR residue had at least one heavy atom within 4 Å of a binder heavy atom in the complex prediction. Mapping the contact frequencies from all submissions onto a structure of EGFR‘s extracellular region illustrates that the designs strongly favored sites in Domains I and III, with little to no utilization of Domains II and IV (Figure 19A).

**Figure 19:**
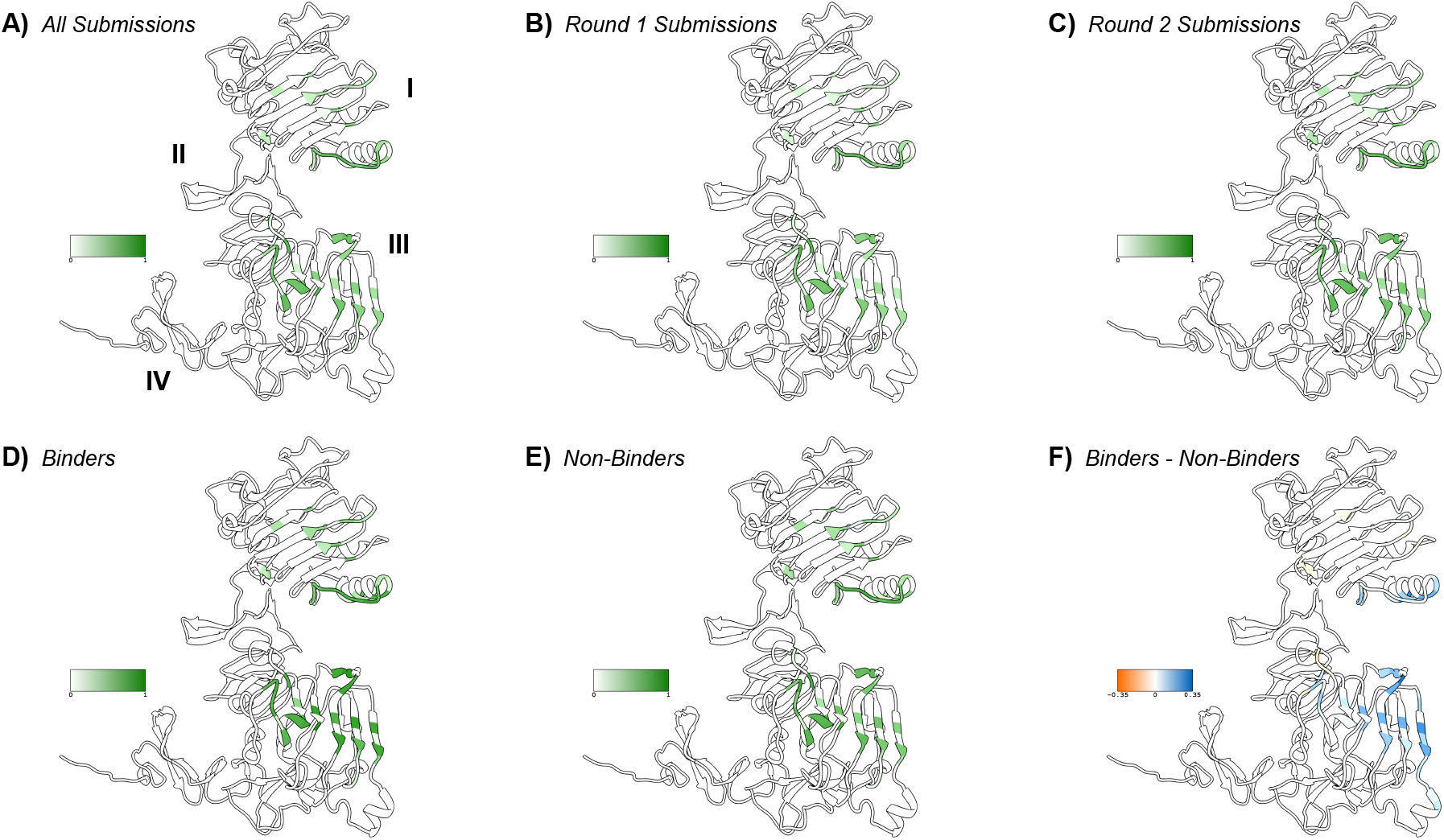
Contact frequencies mapped onto a structure of EGFR for (A) all submitted designs from both rounds, (B) submissions from Round 1, (C) submissions from Round 2, (D) designs which successfully bound when tested in the lab, and designs that did not show binding when tested. (F) Structural visualization of the contact frequency differences between successful binders and non-binders that were tested experimentally.

There are various possible reasons for this preference. The leucine-rich or L domains (I and III) include binding sites both for natural ligands and for FDA-approved therapeutic antibodies [Ogi+02; Gar+02; Li+05; Voi+12; LKF08]. Furthermore, L domains from EGFR, Insulin-like Growth Factor 1 Receptor (IGF1R), and the insulin receptor have been used as targets in recent *de novo* binder design studies [Cao+22; Wat+23] that preceded this competition. EGFR‘s domains of this type were therefore similar to examples in the literature that designers may have been familiar with. Social factors were also relevant to designers’ binding site choices; for instance, target truncation strategies were discussed on X [Bri24]. Importantly, AlphaFold’s prediction capabilities are an additional consideration affecting our analysis of contacts: Predicted contacts do not necessarily represent either true contacts or the sites designers intended to target, and it is possible that AlphaFold struggled to predict contacts in some EGFR regions more than others, or for some types of binders more than others.

The preference for Domains I and III was clear in both rounds of the competition (Figure 19B and C), even though only R2 had specified that binders should compete with EGF. The R2 submissions appeared to show slightly greater contact frequencies compared to R1 for some residues in Domain I and for outward-facing residues on the relatively flat surface of Domain III. The latter region encompasses residues S440 and G441, which were among the recommended hotspots in R2.

Overall, successful binders and non-binders targeted the same locations on EGFR, but with different relative frequencies (Figure 19D and E). To visualize which target contacts were enriched among successful binders, we mapped the contact frequency differences between successful binders and non-binders onto the structure of EGFR. The N-terminal portion of Domain I and the flat region of Domain III displayed the most substantial enrichment (Figure 19F), with the largest positive differences in contact frequency (> 0.25) at residues Q411, R29, and K465. Interestingly, these EGFR residues are hydrophilic, despite previous work indicating that hydrophobic sites yield more favorable outcomes in computational binder design campaigns [Cao+22].

To explore whether this enrichment reflected the higher success rate of designs derived from known binders compared to *de novo* designs, we decomposed the contact frequency differences between binders and non-binders (Figure 20A) based on design method. Among the designs with known binding outcomes (i.e., those that expressed), residues in both Domain I and Domain III showed contact frequency differences between designs labeled as “optimized” or “diversified” and those labeled as “*de novo*” or “hallucination” (Figure 20B). The pattern of contact enrichment in optimized/diversified designs partially mirrors the set of EGFR residue contacts overrepresented in successful binders, but not perfectly: There are Domain I residues farther from the N-terminus that are notably enriched only in the former, suggesting that the effectiveness of different method categories may not fully explain the contact frequency disparities between successful and unsuccessful designs. Meanwhile within the expressed optimized/diversified designs, the contact frequency differences are within a smaller range (Figure 20C), indicating less distinction in contacts between binding and non-binding designs among those based on existing binders. In contrast, binders from *de novo* design or hallucination exhibited more drastic contact frequency differences (Figure 20D). While the two competition rounds produced only 10 experimentally validated binders from these methods, these binders displayed a preference for sites in Domain III and only the N-terminal part of Domain I, with other contacts, mostly later in Domain I, instead depleted among successful binders (Figure 20D and E).

**Figure 20:**
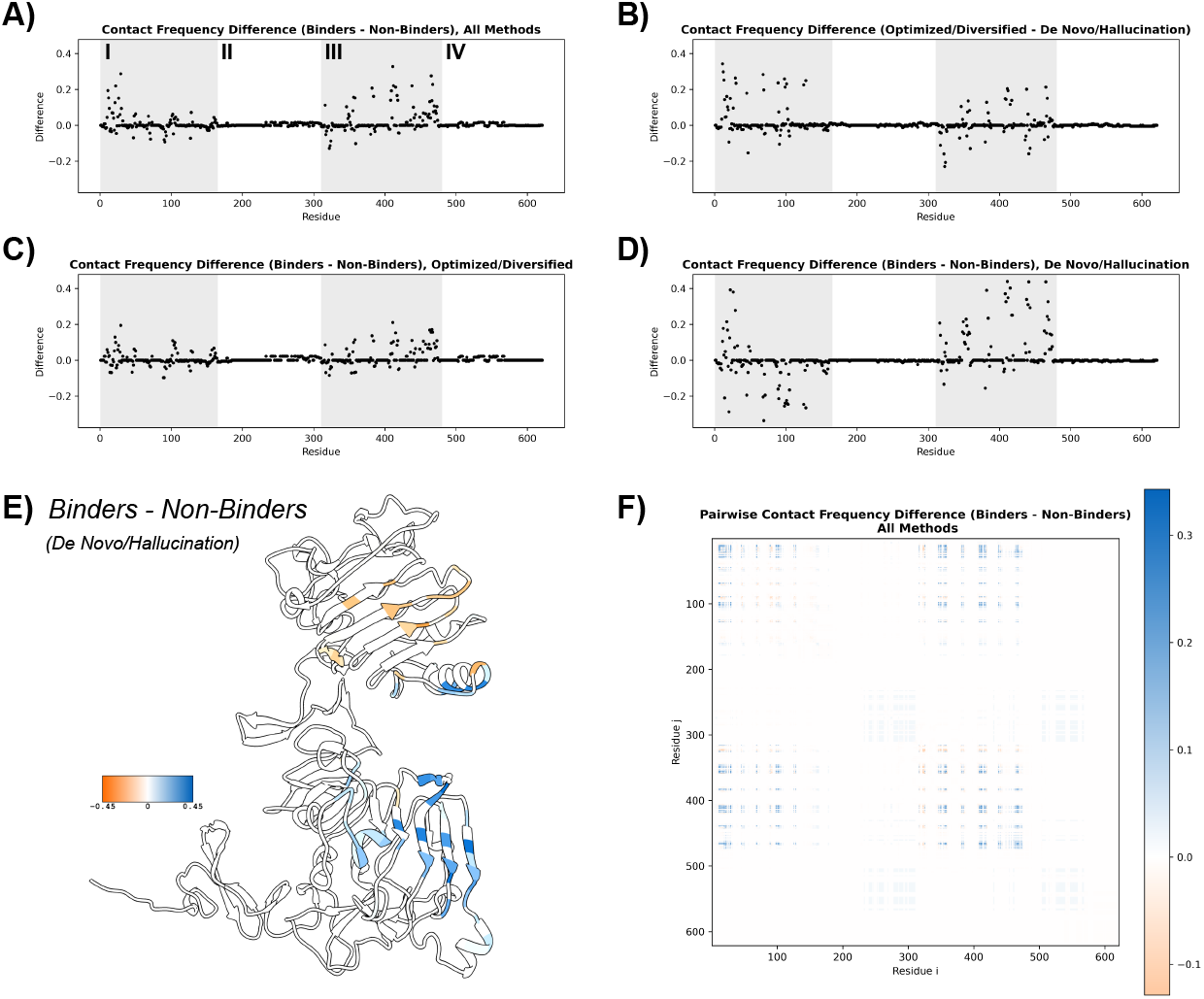
Contact frequency differences by EGFR residue (A) between successful binders and non-binders from all methods, (B) between expressed designs labeled as optimized or diversified and expressed designs from *de novo* or hallucination methods, (C) between successful binders and non-binders among optimized/diversified designs, and (D) between successful binders and non-binders among *de novo*/hallucinated designs. (E) Structural visualization of the contact frequency differences between successful binders and non-binders among *de novo*/hallucinated designs. (F) Pairwise contact frequency differences between successful binders and non-binders from all methods.

Lastly, considering the apparent prevalence of contacts in Domain III and the N-terminal portion of Domain I among validated binders, we sought to determine whether binders frequently contacted these two regions simultaneously or instead contacted only one or the other. We therefore examined pairwise contact frequencies, or the fraction of predicted structures in which two specific EGFR residues were both in contact with the binder, for successful binders and non-binders from all methods. The pairwise contact frequency differences showed off-diagonal enrichment in simultaneous Domain I and Domain III contacts among successful binders (Figure 20F), demonstrating a greater representation of designs which, like the endogenous ligands EGF and TGF-α, interact with both L domains.

### A.6 Description of ANOVA metrics

The AlphaPulldown pipeline provides PTM, PAE, and pLDDT values using AlphaFold2, mpDockQ-pDockQ scores using the formula provided by its authors, and physiochemical and geometrical properties from the PI-score pipeline.

- *Num_intf_residues*: number of residues at the interface.
- *Polar*: number of polar residues (Ser, Thr, Asn, Gln, His, and Tyr) at the interface.
- *Hydrophobic*: number of hydrophobic residues (Ala, Leu, Ile, Val phe, Trp, Cys, Met) at the interface.
- *Charged*: number of charged residues (Asp, Glu, Lys, Arg) at the interface.
- *contact_pairs*: number of atomic contacts between the interface residues.
- *sc*: geometric shape complementarity of protein-protein interfaces. sc ranges between 0 and 1, with sc = 1 meaning two proteins mesh precisely.
- *hb*: number of hydrogen bonds in the interface.
- *sb*: number of salt bridges at the interface.
- *int_solv_en*: interface solvation energy.
- *int_area*: interface surface area that will be inaccessible to solvent upon interface formation.

## Footnotes

1 https://github.com/adaptyvbio/competition_metrics

2 e.g. peptides, minibinders, nanobodies, antibodies

## References

[Kuh+03] Brian Kuhlman et al. “Design of a Novel Globular Protein Fold with Atomic-Level Accuracy”. In: Science 302.5649 (2003). Publisher: American Association for the Advancement of Science, pp. 1364–1368. doi: 10.1126/science.1089427. url: https://www.science.org/doi/10.1126/science.1089427 (visited on 02/07/2025) (cit. on p. 2).

[Fle+11] Sarel J Fleishman et al. “Computational design of proteins targeting the conserved stem region of influenza hemagglutinin”. In: Science 332.6031 (2011), pp. 816–821 (cit. on p. 2).

[Jum+21] John Jumper et al. “Highly accurate protein structure prediction with AlphaFold”. In: Nature 596.7873 (2021), pp. 583–589. issn: 1476-4687. doi: 10.1038/s41586-021-03819-2. url: https://doi.org/10.1038/s41586-021-03819-2 (cit. on pp. 2, 3, 5).

[Eva+22] Richard Evans et al. “Protein complex prediction with AlphaFold-Multimer”. In: bioRxiv (2022). doi: 10.1101/2021.10.04.463034. eprint: https://www.biorxiv.org/content/early/2022/03/10/2021.10.04.463034.full.pdf. url: https://www.biorxiv.org/content/early/2022/03/10/2021.10.04.463034 (cit. on pp. 2, 3, 5).

[Ben+23] Nathaniel R. Bennett et al. “Improving de novo protein binder design with deep learning”. In: Nature Communications 2023 14:1 14 (1 2023), pp. 1–9. issn: 2041-1723. doi: 10.1038/s41467-023-38328-5. url: https://www.nature.com/articles/s41467-023-38328-5 (cit. on pp. 2, 3, 11).

[CLH24] Alexander E Chu, Tianyu Lu, and Po-Ssu Huang. “Sparks of function by de novo protein design”. In: Nature biotechnology 42.2 (2024), pp. 203–215 (cit. on p. 2).

[Che+17] Aaron Chevalier et al. “Massively parallel de novo protein design for targeted therapeutics”. In: Nature 2017 550:7674 550 (7674 Sept. 2017), pp. 74–79. issn: 1476-4687. doi: 10.1038/nature23912. url: https://www.nature.com/articles/nature23912 (cit. on p. 2).

[Cao+20] Longxing Cao et al. “De novo design of picomolar SARS-CoV-2 miniprotein inhibitors”. In: Science 370 (6515 Oct. 2020). issn: 10959203. doi: 10.1126/SCIENCE.ABD9909/SUPPL_FILE/ABD9909-CAO-SM.PDF. url: https://www.science.org/doi/10.1126/science.abd9909 (cit. on p. 2).

[Joh+24] Kristoffer Haurum Johansen et al. “De novo designed pMHC binders facilitate T cell induced killing of cancer cells”. In: bioRxiv (Dec. 2024), p. 2024.11.27.624796. doi: 10.1101/2024.11.27.624796. url: https://www.biorxiv.org/content/10.1101/2024.11.27.624796v1 %20 https://www.biorxiv.org/content/10.1101/2024.11.27.624796v1.abstract (cit. on p. 2).

[Tor+25] Susana Vázquez Torres et al. “De novo designed proteins neutralize lethal snake venom toxins”. In: Nature 2025 639:8053 639 (8053 Jan. 2025), pp. 225–231. issn: 1476-4687. doi: 10.1038/s41586-024-08393-x. url: https://www.nature.com/articles/s41586-024-08393-x (cit. on p. 2).

[Cao+22] Longxing Cao et al. “Design of protein-binding proteins from the target structure alone”. In: Nature 605.7910 (2022), pp. 551–560 (cit. on pp. 2, 4, 14, 22, 25, 26).

[Wat+23] Joseph L. Watson et al. “De novo design of protein structure and function with RFdiffusion”. In: Nature 620.7976 (2023), pp. 1089–1100. issn: 1476-4687. doi: 10.1038/s41586-023-06415-8. url: https://doi.org/10.1038/s41586-023-06415-8 (cit. on pp. 2, 4, 6, 7, 10, 14, 25).

[Arm+24] Chase Armer et al. “Results of the Protein Engineering Tournament: An Open Science Benchmark for Protein Modeling and Design”. In: bioRxiv (Aug. 12, 2024), p. 2024.08.12.606135. doi: 10.1101/2024.08.12.606135. url: https://www.biorxiv.org/content/10.1101/2024.08.12.606135 (cit. on pp. 2, 13).

[Era+24] M. Frank Erasmus et al. “AIntibody: an experimentally validated in silico antibody discovery design challenge”. In: Nature Biotechnology 2024 42:11 42 (11 2024), pp. 1637–1642. issn: 1546-1696. doi: 10.1038/s41587-024-02469-9. url: https://www.nature.com/articles/s41587-024-02469-9 (cit. on p. 2).

[Tyd+24] Claiborne W. Tydings et al. “Analysis of EGFR binding hotspots for design of new EGFR inhibitory biologics”. In: Protein Science : A Publication of the Protein Society 33 (10 Oct. 2024), e5141. issn: 1469896X. doi: 10.1002/PRO.5141. url: https://pmc.ncbi.nlm.nih.gov/articles/PMC11400634/ (cit. on p. 2).

[UMY21] Mary Luz Uribe, Ilaria Marrocco, and Yosef Yarden. “EGFR in Cancer: Signaling Mechanisms, Drugs, and Acquired Resistance”. In: Cancers 13 (11 June 2021), p. 2748. issn: 20726694. doi: 10.3390/CANCERS13112748. url: https://pmc.ncbi.nlm.nih.gov/articles/PMC8197917/ (cit. on p. 2).

[Don23] David Donoho. Data Science at the Singularity. Oct. 2, 2023. doi: 10.48550/arXiv.2310.00865. arXiv: 2310.00865 [stat]. url: http://arxiv.org/abs/2310.00865 (visited on 01/31/2025). Pre-published (cit. on pp. 2, 15).

[LKF08] Shiqing Li, Paul Kussie, and Kathryn M Ferguson. “Structural basis for EGF receptor inhibition by the therapeutic antibody IMC-11F8”. In: Structure 16.2 (2008), pp. 216–227 (cit. on pp. 2, 25).

[Mir+22] Milot Mirdita et al. “ColabFold: making protein folding accessible to all”. In: Nature Methods 19.6 (2022), pp. 679–682. issn: 1548-7105. doi: 10.1038/s41592-022-01488-1. url: http://dx.doi.org/10.1038/s41592-022-01488-1 (cit. on pp. 3, 6, 7).

[SS17] Martin Steinegger and Johannes Söding. “MMseqs2 enables sensitive protein sequence searching for the analysis of massive data sets”. In: Nat Biotechnol (2017) (cit. on pp. 3, 6).

[Haa24] Nikhil Haas. “Round 1 correlations between metrics and affinity or expression”. In: https://x.com/NikhilHaas/status/1844446275967779284 (2024) (cit. on p. 3).

[Lin+23] Zeming Lin et al. “Evolutionary-scale prediction of atomic-level protein structure with a language model”. In: Science 379.6637 (2023). Publisher: American Association for the Advancement of Science, pp. 1123–1130. doi: 10.1126/science.ade2574. url: https://www.science.org/doi/10.1126/science.ade2574 (visited on 05/31/2024) (cit. on pp. 3, 5, 7, 13).

[Tea24a] Adaptyv Team. “Adaptyv - Docs”. In: https://docs.adaptyvbio.com (2024) (cit. on p. 4).

[Tea24b] Adaptyv Team. “Case Study: Benchmarking RFdiffusion generated binders for IL-7Ra”. In: https://www.adaptyvbio.com/blog/rfdiff_il7ra (2024) (cit. on p. 4).

[CK24a] Tudor-Stefan Cotet and Igor Krawczuk. “Protein Optimization 102: Lessons from the protein design competition”. In: https://www.adaptyvbio.com/blog/po102 (2024) (cit. on pp. 4, 7).

[CK24b] Tudor-Stefan Cotet and Igor Krawczuk. “Protein Optimization 103: Racing to the Top 100”. In: https://www.adaptyvbio.com/blog/po103 (2024) (cit. on pp. 4, 7).

[CK24c] Tudor-Stefan Cotet and Igor Krawczuk. “Protein Optimization 101: Insights from the literature”. In: https://www.adaptyvbio.com/blog/po101 (2024) (cit. on p. 4).

[CK24d] Tudor-Stefan Cotet and Igor Krawczuk. “Protein Design Competition: Has binder design been solved?” In: https://www.adaptyvbio.com/blog/po104 (2024) (cit. on pp. 4, 7).

[Nau25] Brian Naughton. Boolean Biotech: The Adaptyv binder design competition. Sat 07 September 2024. Apr. 2025. url: https://web.archive.org/web/20250415112245/ https://blog.booleanbiotech.com/adaptyv-binder-design-competition#expand (visited on 04/15/2025) (cit. on p. 4).

[Cam25] Max Campbell. Blog: Adaptyv Bio Round 2 - EGFR Binder Design. 6 December 2024. Apr. 2025. url: https://web.archive.org/web/20250415112343/ https://www.maxc.codes/posts/adaptyv-bio-egfr-round-2/ (visited on 04/15/2025) (cit. on p. 4).

[Viz25] Vizuro team. Vizuro Selected for Wet Lab Validation in Winter’24 Adaptyv Bio’s EGFR Binder Design. Jan 10. Apr. 2025. url: https://web.archive.org/web/20250415112455/ https://www.vizuro.com/post/vizuro-selected-for-wet-lab-validation-in-winter24 https://www.vizuro.com/post/vizuro-selected-for-wet-lab-validation-in-winter24-adaptyv-bio-s-egfr-binder-design (visited on 04/15/2025) (cit. on p. 4).

[Ale24] Alex Naka [gottapatchemall]. 1/I did my own little hackathon last weekend designing EGFR binders for adaptyvbio’s protein design competition. I was really excited to see that my submissions took the top 10 spots in the virtual scoring phase! I got some DMs asking about my process so here’s a thread: https://t.co/6lypapUywJ. en. Tweet. Aug. 2024. url: https://x.com/gottapatchemall/status/1827386015713260019 (visited on 04/15/2025) (cit. on p. 4).

[Nik24] Nikhil Haas [NikhilHaas]. adaptyvbio we put together stats on what correlates with expression and binding for 248 seqs from the EGFR Competition + data from the exp we ran w/you. binding energy and a couple other scores might work across various seq lengths for R2 eval criteria https://t.co/AiCoWfSK8I. en. Tweet. Oct. 2024. url: https://x.com/NikhilHaas/status/1844446275967779284 (visited on 04/15/2025) (cit. on p. 4).

[Git25] Anthony Gitter. agitter/adaptyvbio-egfr: Sequences from Adaptyv Bio’s EGFR Protein Design Competition. Accessed 2025-04-15. Mar. 2025. doi: 10.5281/zenodo.13972594. url: https://github.com/agitter/adaptyvbio-egfr (visited on 04/15/2025) (cit. on p. 4).

[Suz25] Shosuke Suzuki. suzuki-2001/adaptyv-protein-comp: Design data and process for the AdaptyvBio protein design competition. Accessed 2025-04-15. Mar. 2025. url: https://github.com/suzuki-2001/adaptyv-protein-comp (visited on 04/15/2025) (cit. on p. 4).

[Fer25] Constance Ferragu. “Optimizing Cetuximab”. In: https://www.cradle.bio/blog/adaptyv2 (2025) (cit. on p. 4).

[Cal24] Ewen Callaway. “AI has dreamt up a blizzard of new proteins. Do any of them actually work?” en. In: Nature 634.8034 (Oct. 2024), pp. 532–533. doi: 10.1038/d41586-024-03335-z. url: https://www.nature.com/articles/d41586-024-03335-z (visited on 04/15/2025) (cit. on p. 5).

[Sto+24] Filippo Stocco et al. “Guiding Generative Protein Language Models with Reinforcement Learning”. In: (Dec. 2024). url: https://arxiv.org/abs/2412.12979v2 (cit. on pp. 5, 7).

[Pac+24] Martin Pacesa et al. “BindCraft: one-shot design of functional protein binders”. Version 1. In: bioRxiv (Dec. 7, 2024), p. 2024.09.30.615802. doi: url{10.1101/2024.09.30.615802}. url: %5Curl%7B https://www.biorxiv.org/content/10.1101/2024.09.30.615802v2%7D (visited on 01/31/2025) (cit. on pp. 5, 10, 14, 15, 22, 23).

[Gov+24] Casper A. Goverde et al. “Computational design of soluble and functional membrane protein analogues”. In: Nature 631.8020 (2024), pp. 449–458. issn: 1476-4687. doi: 10.1038/s41586-024-07601-y. url: https://doi.org/10.1038/s41586-024-07601-y (cit. on pp. 5, 7, 10, 15).

[Dau+22] J. Dauparas et al. “Robust deep learning–based protein sequence design using ProteinMPNN”. In: Science 378.6615 (Oct. 2022). Publisher: American Association for the Advancement of Science, pp. 49–56. doi: 10.1126/science.add2187. url: https://www.science.org/doi/10.1126/science.add2187 (visited on 02/04/2023) (cit. on pp. 5, 6, 9).

[Alf+17] Rebecca F Alford et al. “The Rosetta all-atom energy function for macromolecular modeling and design”. In: Journal of chemical theory and computation 13.6 (2017), pp. 3031–3048 (cit. on pp. 5, 7).

[Hut+97] Jeffrey R Huth et al. “Design of an expression system for detecting folded protein domains and mapping macromolecular interactions by NMR”. In: Protein science 6.11 (1997), pp. 2359–2364 (cit. on p. 5).

[Gel+25] Sam Gelman et al. “Biophysics-based protein language models for protein engineering”. en. In: bioRxiv (2025), p. 2024.03.15.585128. doi: 10.1101/2024.03.15.585128. url: https://www.biorxiv.org/content/10.1101/2024.03.15.585128v2 (visited on 02/07/2025) (cit. on p. 5).

[Fer+24] Constance Ferragu et al. “8x improvement in EGFR binding affinity: winning the Adaptyv Bio protein design competition”. In: https://www.cradle.bio/blog/adaptyv-protein-design-competition (2024) (cit. on pp. 5, 10).

[Zho+20] Jeffrey O Zhou et al. “The Effects of Framework Mutations at the Variable Domain Interface on Antibody Affinity Maturation in an HIV-1 Broadly Neutralizing Antibody Lineage”. In: Front Immunol. (2020) (cit. on p. 5).

[Cas+24] Leonardo V Castorina et al. “TIMED-Design: flexible and accessible protein sequence design with convolutional neural networks”. In: Protein Engineering, Design and Selection 37 (2024). issn: 1741-0134. doi: 10.1093/protein/gzae002. url: http://dx.doi.org/10.1093/protein/gzae002 (cit. on pp. 6, 9).

[SW21] Michael J Stam and Christopher W Wood. “DE-STRESS: a user-friendly web application for the evaluation of protein designs”. In: Protein Engineering, Design and Selection 34 (2021). issn: 1741-0134. doi: 10.1093/protein/gzab029. url: http://dx.doi.org/10.1093/protein/gzab029 (cit. on p. 6).

[Thu+21] Vineet Thumuluri et al. “NetSolP: predicting protein solubility in Escherichia coli using language models”. In: Bioinformatics 38.4 (2021). Ed. by Alfonso Valencia, pp. 941–946. issn: 1367-4811. doi: 10.1093/bioinformatics/btab801. url: http://dx.doi.org/10.1093/bioinformatics/btab801 (cit. on pp. 6, 7).

[Riv+21] Alexander Rives et al. “Biological structure and function emerge from scaling unsupervised learning to 250 million protein sequences”. In: Proceedings of the National Academy of Sciences of the United States of America 118 (15 Apr. 2021), e2016239118. issn: 10916490. doi: 10.1073/PNAS.2016239118/SUPPL_FILE/PNAS.2016239118.SAPP.PDF. url: https://www.pnas.org/doi/abs/10.1073/pnas.2016239118 (cit. on pp. 6, 7).

[Kur+19] Aleksander Kuriata et al. “Aggrescan3D (A3D) 2.0: prediction and engineering of protein solubility”. In: Nucleic Acids Research 47.W1 (2019), W300–W307. issn: 1362-4962. doi: 10.1093/nar/gkz321. url: http://dx.doi.org/10.1093/nar/gkz321 (cit. on p. 6).

[QZ20] Yifei Qi and John Z. H. Zhang. “DenseCPD: Improving the Accuracy of Neural-Network-Based Computational Protein Sequence Design with DenseNet”. In: Journal of Chemical Information and Modeling 60.3 (2020), pp. 1245–1252. issn: 1549-960X. doi: 10.1021/acs.jcim.0c00043. url: http://dx.doi.org/10.1021/acs.jcim.0c00043 (cit. on p. 6).

[McI+14] Simon McIntosh-Smith et al. “High performance in silico virtual drug screening on manycore processors”. In: The International Journal of High Performance Computing Applications 29.2 (2014), pp. 119–134. issn: 1741-2846. doi: 10.1177/1094342014528252. url: http://dx.doi.org/10.1177/1094342014528252 (cit. on p. 6).

[SNW24] Eugene Shrimpton-Phoenix, Evangelia Notari, and Christopher W. Wood. “drMD: Molecular Dynamics for Experimentalists”. In: (2024). doi: 10.1101/2024.10.29.620839. url: http://dx.doi.org/10.1101/2024.10.29.620839 (cit. on p. 6).

[Bae+21] Minkyung Baek et al. “Accurate prediction of protein structures and interactions using a three-track neural network”. In: Science 373.6557 (2021), pp. 871–876 (cit. on p. 6).

[KH07] Evgeny Krissinel and Kim Henrick. “Inference of macromolecular assemblies from crystalline state”. In: Journal of molecular biology 372.3 (2007), pp. 774–797 (cit. on p. 7).

[Dau+23] Justas Dauparas et al. “Atomic context-conditioned protein sequence design using LigandMPNN”. In: Biorxiv (2023), pp. 2023–12 (cit. on p. 7).

[Mad+23] Ali Madani et al. “Large language models generate functional protein sequences across diverse families”. In: Nature Biotechnology 2023 41:8 41 (8 Jan. 2023), pp. 1099–1106. issn: 1546-1696. doi: 10.1038/s41587-022-01618-2. url: https://www.nature.com/articles/s41587-022-01618-2 (cit. on p. 7).

[FSH22] Noelia Ferruz, Steffen Schmidt, and Birte Höcker. “ProtGPT2 is a deep unsupervised language model for protein design”. In: Nature Communications 2022 13:1 13 (1 July 2022), pp. 1–10. issn: 2041-1723. doi: 10.1038/s41467-022-32007-7. url: https://www.nature.com/articles/s41467-022-32007-7 (cit. on p. 7).

[Eln+20] Ahmed Elnaggar et al. “ProtTrans: Towards Cracking the Language of Life’s Code Through Self-Supervised Deep Learning and High Performance Computing”. In: bioRxiv 14 (8 July 2020). issn: 26928205. url: https://arxiv.org/abs/2007.06225v3 (cit. on p. 7).

[Hay+25] Thomas Hayes et al. “Simulating 500 million years of evolution with a language model”. en. In: Science 387 (6736 Feb. 2025). Pages: 2024.07.01.600583 Section: New Results, pp. 850–858. issn: 0036-8075. doi: 10.1126/SCIENCE.ADS0018. url: https://www.science.org/doi/10.1126/science.ads0018 (visited on 02/12/2025) (cit. on pp. 7, 11).

[RGS21] Jeffrey A. Ruffolo, Jeffrey J. Gray, and Jeremias Sulam. “Deciphering antibody affinity maturation with language models and weakly supervised learning”. In: (Dec. 2021). url: https://arxiv.org/abs/2112.07782v1 (cit. on p. 7).

[RSG22] Jeffrey A. Ruffolo, Jeremias Sulam, and Jeffrey J. Gray. “Antibody structure prediction using interpretable deep learning”. In: Patterns 3 (2 Feb. 2022), p. 100406. issn: 2666-3899. doi: 10.1016/J.PATTER.2021.100406 (cit. on p. 7).

[Mun+24] Geraldene Munsamy et al. “Conditional language models enable the efficient design of proficient enzymes”. In: bioRxiv (May 2024), p. 2024.05.03.592223. doi: 10.1101/2024.05.03.592223. url: https://www.biorxiv.org/content/10.1101/2024.05.03.592223v2 %20 https://www.biorxiv.org/content/10.1101/2024.05.03.592223v2.abstract (cit. on p. 7).

[Bio25] Adaptyv Bio. EGFR2024 Post-Competition Data and Script Collection. Accessed: 2025-04-14. 2025. url: https://github.com/adaptyvbio/egfr2024_post_competition (cit. on p. 7).

[Hon+21] Jiri Hon et al. “SoluProt: prediction of soluble protein expression in Escherichia coli”. In: Bioinformatics 37 (1 2021), pp. 23–28. issn: 1367-4803. doi: \url{10.1093/BIOINFORMATICS/BTAA1102}. url: %5Curl%7B https://dx.doi.org/10.1093/bioinformatics/btaa1102%7D (cit. on p. 7).

[Kem+23] Michel van Kempen et al. “Fast and accurate protein structure search with Foldseek”. In: Nature Biotechnology 2023 42:2 42 (2 May 2023), pp. 243–246. issn: 1546-1696. doi: 10.1038/s41587-023-01773-0. url: https://www.nature.com/articles/s41587-023-01773-0 (cit. on p. 8).

[Hie+23] Brian L. Hie et al. “Efficient evolution of human antibodies from general protein language models”. In: Nature Biotechnology 2023 42:2 42 (2 2023), pp. 275–283. issn: 1546-1696. doi: \url{10.1038/s41587-023-01763-2}. url: https://www.nature.com/articles/s41587-023-01763-2 (cit. on p. 10).

[Sha+24] Varun R. Shanker et al. “Unsupervised evolution of protein and antibody complexes with a structure-informed language model”. In: Science (New York, N.Y.) 385 (6704 2024), pp. 46–53. issn: 10959203. doi: \url{10.1126/SCIENCE.ADK8946/SUPPL_FILE/SCIENCE.ADK8946_MDAR_REPRODUCIBILITY_CHECKLIST.PDF}. url: %5Curl%7B https://www.science.org/doi/10.1126/science.adk8946%7D (cit. on p. 10).

[Ani+21] Ivan Anishchenko et al. “De novo protein design by deep network hallucination”. In: Nature 2021 600:7889 600 (7889 2021), pp. 547–552. issn: 1476-4687. doi: 10.1038/s41586-021-04184-w. url: https://www.nature.com/articles/s41586-021-04184-w (cit. on p. 10).

[Wic+22] B. I.M. Wicky et al. “Hallucinating symmetric protein assemblies”. In: Science 378 (6615 2022), p. 2025. issn: 10959203. doi: \url{10.1126/SCIENCE.ADD1964/SUPPL_FILE/SCIENCE.ADD1964_SM.PDF}. url: https://www.science.org/doi/10.1126/science.add1964 (cit. on p. 10).

[Fra+24] Christopher Frank et al. “Scalable protein design using optimization in a relaxed sequence space”. In: Science (New York, N.Y.) 386 (6720 2024), pp. 439–445. issn: 10959203. doi: \url{10.1126/SCIENCE.ADQ1741/SUPPL_FILE/SCIENCE.ADQ1741_MDAR_REPRODUCIBILITY_CHECKLIST.PDF}. url: %5Curl% 7B https://www.science.org/doi/10.1126/science.adq1741%7D (cit. on p. 10).

[UMG24a] Talip Uçar, Cedric Malherbe, and Ferran Gonzalez. “Exploring Log-Likelihood Scores for Ranking Antibody Sequence Designs”. In: bioRxiv (Oct. 2024), p. 2024.10.07.617023. doi: 10.1101/2024.10.07.617023. url: https://www.biorxiv.org/content/10.1101/2024.10.07.617023v4 %20 https://www.biorxiv.org/content/10.1101/2024.10.07.617023v4.abstract (cit. on p. 11).

[CRG24] Michael Chungyoun, Jeffrey Ruffolo, and Jeffrey Gray. “FLAb: Benchmarking deep learning methods for antibody fitness prediction”. en. In: bioRxiv (2024). doi: 10.1101/2024.01.13.575504. eprint: https://www.biorxiv.org/content/early/2024/01/15/2024.01.13.575504.full.pdf. url: https://www.biorxiv.org/content/10.1101/2024.01.13.575504v1 (visited on 02/12/2025) (cit. on pp. 11–13).

[ESM24] ESM Team. ESM Cambrian: Revealing the mysteries of proteins with unsupervised learning. EvolutionaryScale Website. Dec. 2024. url: https://evolutionaryscale.ai/blog/esm-cambrian (visited on 04/15/2025) (cit. on p. 11).

[OBD22] Tobias H. Olsen, Fergus Boyles, and Charlotte M. Deane. “Observed Antibody Space: A diverse database of cleaned, annotated, and translated unpaired and paired antibody sequences”. In: Protein Science 31 (1 Jan. 2022), pp. 141–146. issn: 1469-896X. doi: 10.1002/PRO.4205. url: https://onlinelibrary.wiley.com/doi/full/10.1002/pro.4205 %20 https://onlinelibrary.wiley.com/doi/abs/10.1002/pro.4205 %20 https://onlinelibrary.wiley.com/doi/10.1002/pro.4205 (cit. on p. 11).

[Ada23] Adaptyv Bio. EGFR Competition 1. https://github.com/adaptyvbio/egfr_competition_1. GitHub repository. Accessed: 2025-01-08. 2023 (cit. on p. 11).

[WSK24] Jordan R. Willis, Troy M. Sincomb, and Caleb K. Kibet. sadie-antibody: Structural Analysis and Design Interface for Engineering antibodies. https://pypi.org/project/sadie-antibody/. Python package. Accessed: 2025-04-08. 2024 (cit. on p. 11).

[Bio24] BioLM. Silica: BioML Hackathon Team Repository. Accessed: 2025-04-11. 2024. url: https://github.com/BioLM/silica (cit. on p. 11).

[Lux] Lux Capital and Evolutionary Scale and Enveda Biosciences. Bio x ML Hackathon 2024. https://hackathon.bio/2024. Held online from mOctober 10–20, 2024. (Visited on 04/15/2025) (cit. on p. 11).

[Aba+23] Brennan Abanades et al. “ImmuneBuilder: Deep-Learning models for predicting the structures of immune proteins”. en. In: Communications Biology 6.1 (2023). Publisher: Nature Publishing Group, pp. 1–8. issn: 2399-3642. doi: 10.1038/s42003-023-04927-7. url: https://www.nature.com/articles/s42003-023-04927-7 (visited on 02/12/2025) (cit. on p. 12).

[Ken+24] Henry Kenlay et al. “ABodyBuilder3: improved and scalable antibody structure predictions”. In: Bioinformatics 40.10 (Oct. 2024), btae576. doi: 10.1093/bioinformatics/btae576. url: https://academic.oup.com/bioinformatics/article/40/10/btae576/7810444 (cit. on p. 12).

[UMG24b] Talip Ucar, Cedric Malherbe, and Ferran Gonzalez. “Exploring Log-Likelihood Scores for Ranking Antibody Sequence Designs”. Version 4. In: bioRxiv (2024). doi: \url{10.1101/2024.10.07.617023}. url: %5Curl%7B%20 https://www.biorxiv.org/content/10.1101/2024.10.07.617023v4.full.pdf%7D (cit. on p. 12).

[EG25] Floriane Eshak and Anne Goupil-Lamy. “Advancements in Nanobody Epitope Prediction: A Comparative Study of AlphaFold2Multimer vs AlphaFold3”. In: Journal of Chemical Information and Modeling 65.4 (2025), pp. 1234–1245. doi: 10.1021/acs.jcim.4c01877. url: https://pubs.acs.org/doi/10.1021/acs.jcim.4c01877 (cit. on p. 12).

[Bry+22] Patrick Bryant et al. “Predicting the structure of large protein complexes using AlphaFold and Monte Carlo tree search”. In: Nature Communications 13.1 (Oct. 2022), p. 6028. doi: 10.1038/s41467-022-33729-4. url: https://doi.org/10.1038/s41467-022-33729-4 (visited on 04/15/2025) (cit. on p. 12).

[Mal] Sony Malhotra. ppi_scoring Wiki. https://gitlab.com/sm2185/ppi_scoring/-/wikis/home. Accessed: 2025-04-15 (cit. on p. 12).

[Yu+23] Dingquan Yu et al. “AlphaPulldown–a Python Package for Protein–Protein Interaction Screens Using AlphaFold-Multimer”. In: Bioinformatics 39.1 (2023), btac749. doi: 10.1093/bioinformatics/btac749. url: https://academic.oup.com/bioinformatics/article/39/1/btac749/6839971 (visited on 04/15/2025) (cit. on p. 12).

[Xue+16] Li C. Xue et al. “PRODIGY: a web server for predicting the binding affinity of protein–protein complexes”. In: Bioinformatics 32 (23 Dec. 2016), pp. 3676–3678. issn: 1367-4803. doi: 10.1093/BIOINFORMATICS/BTW514. url: https://dx.doi.org/10.1093/bioinformatics/btw514 (cit. on p. 13).

[MBB24] Kevin Michalewicz, Mauricio Barahona, and Barbara Bravi. “ANTIPASTI: Interpretable prediction of antibody binding affinity exploiting normal modes and deep learning”. In: Structure 32 (12 Dec. 2024), 2422–2434.e5. issn: 0969-2126. doi: 10.1016/J.STR.2024.10.001. url: https://linkinghub.elsevier.com/retrieve/pii/S0969212624004362 (cit. on p. 13).

[Zam+24] Vinicius Zambaldi et al. “De novo design of high-affinity protein binders with AlphaProteo”. In: ArXiv preprint abs/2409.08022 (2024). url: https://arxiv.org/abs/2409.08022(cit. on p. 14).

[Wan+18] Alex Wang et al. “GLUE: A Multi-Task Benchmark and Analysis Platform for Natural Language Understanding”. In: EMNLP 2018 - 2018 EMNLP Workshop BlackboxNLP: Analyzing and Interpreting Neural Networks for NLP, Proceedings of the 1st Workshop (Apr. 2018), pp. 353–355. doi: 10.18653/v1/w18-5446. url: https://arxiv.org/abs/1804.07461v3 (cit. on p. 14).

[Wan+19] Alex Wang et al. “SuperGLUE: A Stickier Benchmark for General-Purpose Language Understanding Systems”. In: Advances in Neural Information Processing Systems 32 (May 2019). issn: 10495258. url: https://arxiv.org/abs/1905.00537v3 (cit. on p. 14).

[Hen+20] Dan Hendrycks et al. “Measuring Massive Multitask Language Understanding”. In: ICLR 2021 - 9th International Conference on Learning Representations (Sept. 2020). url: https://arxiv.org/abs/2009.03300v3 (cit. on p. 14).

[Den+09] Jia Deng et al. “ImageNet: A Large-Scale Hierarchical Image Database”. In: 2009 IEEE Conference on Computer Vision and Pattern Recognition, CVPR 2009 (2009), pp. 248–255. doi: 10.1109/CVPR.2009.5206848 (cit. on p. 14).

[Ale+24] Reem Aleithan et al. “SWE-Bench+: Enhanced Coding Benchmark for LLMs”. In: (Oct. 2024).url: https://arxiv.org/abs/2410.06992v2 (cit. on p. 14).

[Jim+23] Carlos E. Jimenez et al. “SWE-bench: Can Language Models Resolve Real-World GitHub Issues?” In: 12th International Conference on Learning Representations, ICLR 2024 (Oct. 2023). url: https://arxiv.org/abs/2310.06770v3 (cit. on p. 14).

[Cen06] Center for High Throughput Computing. Center for High Throughput Computing. 2006. doi: 10.21231/GNT1-HW21 (cit. on p. 15).

[Ogi+02] Hideo Ogiso et al. “Crystal structure of the complex of human epidermal growth factor and receptor extracellular domains”. In: Cell 110.6 (2002), pp. 775–787 (cit. on p. 25).

[Gar+02] Thomas PJ Garrett et al. “Crystal structure of a truncated epidermal growth factor receptor extracellular domain bound to transforming growth factor α”. In: Cell 110.6 (2002), pp. 763–773 (cit. on p. 25).

[Li+05] Shiqing Li et al. “Structural basis for inhibition of the epidermal growth factor receptor by cetuximab”. In: Cancer cell 7.4 (2005), pp. 301–311 (cit. on p. 25).

[Voi+12] Mareike Voigt et al. “Functional dissection of the epidermal growth factor receptor epitopes targeted by panitumumab and cetuximab”. In: Neoplasia 14.11 (2012), 1023–IN3 (cit. on p. 25).

[Bri24] Brian Naughton [btnaughton]. I added basic BindCraft (few options exposed!) to https://github.com/hgbrian/biomodals (code from MartinPacesa’s colab) git clone https://github.com/hgbrian/biomodals cd biomodals wget https://raw.githubusercontent.com/martinpacesa/BindCraft/refs/heads/main/example/PDL1.pdb GPU=A100 modal run modal_bindcraft.py –input-pdb PDL1.pdb –number-of-final-designs 1. en. Tweet. Oct. 2024. url: https://x.com/btnaughton/status/1851266190934753742 (visited on 04/15/2025) (cit. on p. 25).

